# Astral microtubule crosslinking safeguards uniform nuclear distribution in the *Drosophila* syncytium

**DOI:** 10.1101/859975

**Authors:** Ojas Deshpande, Jorge de-Carvalho, Diana V. Vieira, Ivo A. Telley

## Abstract

The early insect embryo develops as a multinucleated cell distributing the genome uniformly to the cell cortex. Mechanistic insight for nuclear positioning beyond cytoskeletal requirements is missing. Contemporary hypotheses propose actomyosin driven cytoplasmic movement transporting nuclei, or repulsion of neighbor nuclei driven by microtubule motors. Here, we show that microtubule crosslinking by Feo and Klp3A is essential for nuclear distribution and internuclear distance maintenance in *Drosophila*. Germline knockdown causes irregular, less dense nuclear delivery to the cell cortex and smaller distribution in *ex vivo* embryo explants. A minimal internuclear distance is maintained in explants from control embryos but not from Feo inhibited embryos, following micromanipulation assisted repositioning. A dimerization deficient Feo abolishes nuclear separation in embryo explants while the full-length protein rescues the genetic knockdown. We conclude that Feo and Klp3A crosslinking of antiparallel microtubule overlap generates a length-regulated mechanical link between neighboring microtubule asters. Enabled by a novel experimental approach, our study illuminates an essential process of embryonic multicellularity.

## Introduction

The nucleus relocates within the cell boundary in response to cell function (Bone and Starr, 2016; Gundersen and Worman, 2013). Aberrant nuclear positioning has been linked to failure of fundamental processes such as early embryo development, cell differentiation, cell migration, polarity determination and homeostasis (Almonacid et al., 2015; 2019; Levy and Holzbaur, 2008; Minc et al., 2011; Neelam et al., 2015; Starr and Han, 2002). In mononuclear cells, cytoskeletal elements mechanically connect the nucleus to the cell cortex and act as the reference system for positioning (Dassow et al., 2009; Pecreaux et al., 2016). One exception are large eggs in which cytoskeletal links between the nucleus and the distant cell cortex are not achieved (Wühr et al., 2009). Conversely, a multinucleated cell – coenocyte – undergoing nuclear proliferation has to disseminate positional information to each additional nucleus and requires a mechanism that adjusts the distance between neighboring nuclei (Gibeaux et al., 2017; Manhart et al., 2018). The early embryo of *Drosophila melanogaster* is both large and multinucleated but exhibits a surprising positional regularity of hundreds of nuclei perturbed by cycles of meta-synchronous nuclear divisions (Foe and Alberts, 1983). During the first seven cycles, the nuclei spread axially from the anterior to the posterior end of the syncytial embryo and occupy the entire volume of the embryo (Baker et al., 1993). During nuclear cycles 7–9, most nuclei migrate to the embryo cortex, where they undergo additional rounds of division as they are anchored and prepared for cellularization (Lecuit and Wieschaus, 2000). Adequate number of nuclei and their proper positioning at the cortex determines cell size (Callaini et al., 1992) and precision of developmental patterning (Petkova et al., 2019), and is a result of regular distribution of ancestor nuclei during the preceding developmental phase (Kao and Megraw, 2009; Megraw et al., 1999; Vaizel-Ohayon and Schejter, 1999). The mechanisms required for maintaining uniform internuclear distances are not understood.

Drug inhibition and mutagenesis suggest that actomyosin mediated cortical contractions drive cytoplasmic streaming and transport the nuclei predominantly along the longer axis of the embryo (Callaini et al., 1992; Dassow and Schubiger, 1994; Deneke et al., 2019; Hatanaka and Okada, 1991; Royou et al., 2004; Wheatley et al., 1995). However, large-scale transport of cytoplasm does neither explain how a uniform distribution emerges nor how nuclei are kept separate. Conversely, astral microtubules from neighboring nuclei may interact and generate repulsive force by motor binding and sliding antiparallel overlaps (Baker et al., 1993), which is reminiscent of the spindle midzone model explaining spindle elongation during anaphase B (Fu et al., 2009; Khmelinskii et al., 2009; Scholey et al., 2016). The effector Klp61F, a homotetrameric, bipolar Kinesin-5 binds two overlapping microtubules and, when microtubules are antiparallel, slides them outwards reducing microtubule overlap length (Cheerambathur et al., 2013; Reinemann et al., 2017; Tao et al., 2006). Fascetto (Feo) is the *Drosophila* homolog of the Ase1p/Prc1/MAP65 family of homodimeric non-motor MAPs that preferentially binds antiparallel microtubule overlaps (Bieling et al., 2010; Subramanian et al., 2010; Vernì et al., 2004). It accumulates at the spindle midzone from anaphase to telophase upon cyclin B degradation and controls the binding affinity of molecular motors in the spindle midzone (Hu et al., 2012; Khmelinskii et al., 2009; Kwon and Scholey, 2004; Wang et al., 2015; Zhu et al., 2006). One of these motors is Klp3A, a Kinesin-4 homolog, an inhibitor of microtubule dynamics (Bieling et al., 2010; Bringmann et al., 2004; Kwon et al., 2004; Subramanian et al., 2013; Williams et al., 1997). Prc1 and Kinesin-4 are sufficient to form a stable microtubule overlap *in vitro* (Bieling et al., 2010). Kinesin-5 can reduce overlapping, antiparallel microtubules crosslinked by Prc1 *in vitro* (Subramanian et al., 2010), which was proposed to contribute to force balance in the spindle midzone during anaphase B (Scholey et al., 2016).

Here, we investigated whether these three proteins are required for nuclear separation, lending support to an aster–aster interaction model (Baker et al., 1993), which has been reconstituted in *Xenopus* egg extract (Nguyen et al., 2014; 2018). We performed a combination of gene knockdown, micromanipulation and perturbation by exogenous protein addition in embryo explants which enable time-lapse visualization of nuclear and cytoskeletal dynamics previously unachieved.

## Results

### Feo localization confirms antiparallel microtubule overlaps between asters of non-sister nuclei

Molecular crosslinking between astral microtubules of neighboring nuclei during the preblastoderm embryo stage has largely been unexplored due to optical constraints in live imaging. Using an extraction method to generate embryo explants from individual preblastoderm embryos (de-Carvalho et al., 2018) expressing Klp61F::GFP and Feo::mCherry, and injected with Alexa647-labeled Tubulin (Fig. 1A), we visualized the localization of Klp61F and Feo to infer about their binding to spindle microtubules (Fig. 1B). Klp61F::GFP localized at the microtubule-organizing centers, the metaphase spindle and the spindle midzone in anaphase, as described previously for the nuclear divisions at the blastoderm stage (Cheerambathur et al., 2013; 2008; Heck et al., 1993; Sharp et al., 1999; Tao et al., 2006) (Suppl. Video 1). Furthermore, during anaphase B and telophase we observed Klp61F::GFP decorated microtubules intercalating with those from the neighboring aster, raising the possibility of antiparallel alignment of these astral microtubules forming an overlap zone to which Kinesin-5 binds. On the other hand, Feo::mCherry exhibited weak localization to the metaphase spindle but strong localization to the spindle midzone during anaphase B and telophase (Fig. 1B, arrows), as previously described for blastoderm division cycles (Wang et al., 2015). Strikingly, Feo also localized as small foci to the region between the nuclei (Fig. 1B,C, arrowheads), thus reporting the presence of antiparallel microtubule overlaps to which Feo binds with higher affinity than individual microtubules. *In vitro*, microtubule overlaps that are decorated by Feo homologs are length controlled through the stabilization activity of Kinesin-4 (Bieling et al., 2010). Thus, the signal of Feo along microtubule overlaps should have a consistent length for a given concentration or activity of Feo and Klp3A. Thus, we measured the length of Feo::mCherry signal foci during anaphase B (Fig. 1D). Because individual microtubules were not resolved, we measured the orientation of the signal foci in the context of where microtubules are growing and radially emanating from the microtubule-organizing centers (MTOC) at the spindle pole. In anaphase and telophase, the four nuclei emerging from any two neighboring spindles define four MTOCs and, thus, four possible combinations of astral microtubule interaction (Fig. 1C, right). We measured the angle *θ* between the long axes of the signal foci and the closest connecting line between two MTOCs (Fig. 1E). *θ* deviated little from zero, supporting the notion that Feo reports microtubule overlaps along the shortest path between neighboring asters. Altogether, in explants from preblastoderm embryos, the relative position of nuclei and the length of astral microtubules lead to the formation of short antiparallel Feo-decorated microtubule overlaps. Furthermore, as a consequence of Feo crosslinking astral microtubules, a mechanical connection is established that may control the distance between neighboring asters and their associated nuclei.

**Figure 1:**
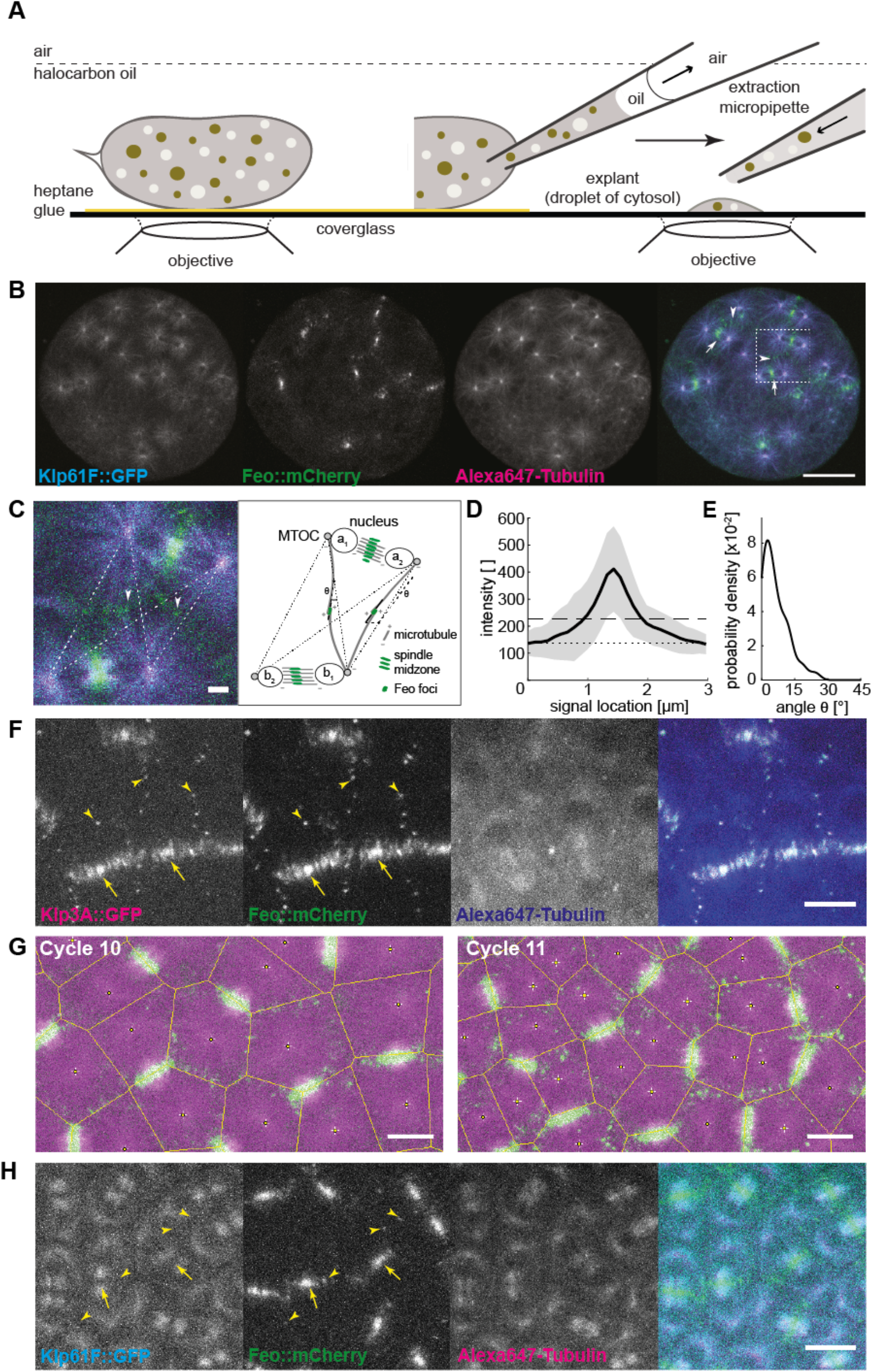
Feo, Klp3A and Klp61F localization confirm antiparallel microtubule overlaps between asters of non-sister nuclei. **A)** Schematic showing a *Drosophila* syncytial embryo immobilized to the coverslip and covered with a thin layer of halocarbon oil ready for time-lapse microscopy. On the right, a preblastoderm embryo is punctured for extraction and deposition of cytosol on the coverslip using a micropipette, thereby generating a series of embryo explants. **B)** Three-color snapshot from a time-lapse (see also Suppl. Video 1) of an explant generated from an embryo expressing Klp61F::GFP (cyan), Feo::mCherry (green) and injected with Alexa647– Tubulin (magenta). During the anaphase/telophase transition, Feo strongly localized to the spindle midzone (arrows) and to the intercalating microtubules from neighboring nuclei (arrowheads). Scale bar, 30 μm. **C)** Zoom-in of the merged color channel image in b) (dashed square). Feo localized as intense foci between neighboring spindles, where microtubules from non-sister nuclei meet (arrowheads show examples). The schematic on the right represents the configuration shown in the image, exemplifying the location of the two pairs of sister nuclei, a^1^–a^2^ and b^1^–b^2^, and two representative Feo foci. The dashed lines represent the shortest path of microtubule interactions between the MTOCs of non-sister nuclei. An intensity profile of the foci was generated by drawing a line (continuous) along the longest axis and centered to the foci. The angle θ relative to the dashed interaction line was determined. Scale bar, 2 μm. **D)** The average intensity profile of Feo foci indicate a foci length of 1.0±0.35 μm. The grey area designates the standard deviation (SD), the dotted line marks the background level, and the dashed line marks two times SD above the background. N=7; n=57. **E)** The distribution of angles (θ) suggests that the antiparallel microtubule overlaps occur mostly along the connecting line between the neighboring non-sister nuclei. N=7; n=42. Cases where foci were symmetric, and a long axis could not be determined were excluded from the analysis. **F)** Three-color snapshot of a blastoderm embryo expressing Klp61F::GFP (cyan), Feo::mCherry (green) and injected with Alexa647–Tubulin (magenta) showed that Feo localized strongly between sister nuclei as part of the spindle midzone (arrows) and, more strikingly, between neighboring non-sister nuclei as distinct foci (arrowheads). Scale bar, 10 μm. Refer to Suppl. Video 2. **G)** Two-color still images of a blastoderm embryo expressing RFP::β-tubulin (magenta) and Feo::GFP (green) during cycles 10 and 11, with Voronoi lines overlaid in yellow. The Voronoi segmentation was calculated with respect to the location of spindle poles and mark all locations with equidistant neighbors. Scale bar, 10 μm. **H)** Three-color snapshot of a blastoderm embryo expressing Klp3A::GFP (magenta), Feo::mCherry (green) and injected with Alexa647–Tubulin (blue) showed that Klp3A co-localizes with Feo at the spindle midzone (arrows) and at the foci between neighboring non-sister nuclei (arrowheads). Scale bar, 10 μm. Refer to Suppl. Video 3.

During the last four syncytial nuclear cycles at the cortex, actin based pseudo-compartments that drive membrane invagination – a physical barrier that is assembled and disassembled in every division cycle (Kotadia et al., 2001; Mavrakis et al., 2009a) – are thought to guarantee nuclear separation. Surprisingly, time-lapse confocal imaging of live embryos expressing Klp3A::GFP and Feo::mCherry (Fig. 1F, Suppl. Video 2), injected with Alexa647–Tubulin, revealed Feo colocalizing with Klp3A at the spindle midzone (arrows) and spot-like signals between neighboring spindles (arrowheads) in anaphase and telophase. On one hand, this observation confirms the combined activity of Feo and Klp3A, whereby Feo binding to microtubule overlaps recruits Klp3A to the overlap (Bieling et al., 2010; Subramanian et al., 2013). On the other hand, the signal foci indicate that antiparallel microtubule overlaps occur between neighboring non-sister nuclei across actin furrows and membrane invaginations. With consecutive division cycles and increasing nuclear packing the foci maintain size but become more frequent (Suppl. Fig. 1A). Triangulation analysis revealed that foci appear at locations equidistant to neighboring spindle poles, in the center of the aster-aster overlap zone (Fig. 1G, Suppl. Fig. 1B), and their size is comparable to those observed in explants (Suppl. Fig. 1C). In embryos expressing Klp61F::GFP and Feo::mCherry the GFP signal is excluded at the spindle midzone where Feo::mCherry localizes strongly. Although the Klp61F concentrates at the spindle poles it does not show distinct localization at the aster-aster interaction zone (Fig. 1H, Suppl. Video 3). Altogether, these observations led us to question the current paradigm that actin pseudo-compartments prevent microtubule crosslinking between neighboring asters or nuclei. We hypothesize from this localization data that the microtubule-based mechanical connection plays a decisive role in nuclear positioning in preblastoderm and early blastoderm stage embryos. Since microtubules are a prerequisite, we perturbed microtubule dynamics with low doses of the depolymerization drug Nocodazole. We supplemented this drug after explant deposition, using a fine micropipette using buffer conditions described elsewhere (Telley et al. 2013). A concentration of 4 μM sustained spindle assembly and chromosomes segregation but abolished daughter nuclei separation (Suppl. Video 4). The microtubule perturbation caused local aggregation of dividing nuclei. We also observed chromosomes fuse and form larger nuclei. Those nuclei more distant from the drug injection site still separated consistently, which we attribute to a diffusion gradient in drug concentration. Overall, this experiment demonstrates that nuclear separation depends on microtubules and supports a role of Feo, Klp3A and Klp61F in nuclear positioning.

### Knockdown of Feo, Klp3A or Klp61F leads to defective nuclear delivery to the embryo cortex

We wanted to understand the functional implication of the three microtubule binding proteins that localize between non-sister nuclei, and if they are required for correct nuclear delivery to the cortex. To this end, we perturbed the protein levels of Feo, Klp3A or Klp61F using an RNA interference (RNAi) approach and UAS–Gal4 expression in the germline (Staller et al., 2013). Using different available (TRiP) fly lines we expressed RNAi against these genes individually in the developing *Drosophila* oocyte (Suppl. Fig. 2A), while expressing Jupiter::GFP, a microtubule reporter (Morin et al., 2001), and H2Av::RFP, a chromatin reporter (Schuh et al., 2007). We exploited the expression kinetics of V32–Gal4 to drive the UASp–RNAi constructs with peak in late oogenesis to prevent undesirable defects during stem cell division. The efficiency of knockdown was measured at the RNA level using a qPCR approach (Table 1, Methods). Fertilization in embryos inhibited for Feo, Klp3A or Klp61F expression was similar to control embryos. However, we were unable to determine the exact cycle number when nuclei arrived at the cortex in knockdown embryos. Of note, the interval of division cycles occurring at the cortex and in the embryo explants remained unaltered when compared to controls. One RNAi construct against *feo* (TRiP # 28926) did not show any phenotype. Under all other knockdown conditions, we observed nuclei arriving later on average; ∼45 min in knockdown condition versus ∼15 min in controls, following a 45 min egg laying period. In knockdown embryos, nuclei were irregularly distributed at the cortex and sometimes missing entirely at the posterior end, in contrast to the regular distribution seen in the control RNAi embryo (Fig. 2A, Suppl. Fig. 2B). The nuclear density is reduced after knockdown as compared to the control (Fig. 2B) but exhibited considerable variability between embryos, which we attributed to variability in UAS-Gal4 mediated expression of RNAi between individual embryos. Reassuringly, in *feo* RNAi embryos we observed a reduction – below detection – of fluorescence intensity of Klp3A::GFP at the midzone and between neighbor asters (Suppl. Fig. 2C). This confirms Klp3A being downstream of Feo binding to microtubule overlaps (Bieling et al., 2010). Our nuclear positioning analysis revealed embryos with larger areas lacking nuclei (Suppl. Fig. 3A), with anatomically eccentric (Suppl. Fig. 3B) and asymmetric nuclear distribution (Suppl. Fig. 3C). Overall, RNAi against *feo* resulted in larger distribution changes than RNAi against *klp61f* (Suppl. Fig. 3) despite similar average internuclear distance (Fig. 2C). The distribution of first neighbor internuclear distance was shifted towards longer distances for knockdown conditions, reflecting larger unoccupied areas, while RNAi against *klp3a* gave on average the strongest phenotype (Fig. 2C, Suppl. Fig. 3). The irregularity in nuclear position at the cortex increased as the cortical nuclear cycles progressed (Suppl. Video 5). Occasionally in *feo* knockdown embryos, we observed sister nuclei fusing after mitosis and non-sister nuclear moving towards each other, leading to fusion of the spindles and over-condensed chromatin. This likely occurs in preblastoderm cycles as well, leading to fewer nuclei arriving at the cortex and contributing to nonuniformity. Conversely, fusion was never seen in controls. In summary, the activity of all three microtubule associated proteins is required in the preblastoderm embryo for correct delivery of nuclei to the embryo cortex. However, Kinesin-5 is required for spindle assembly (Heck et al., 1993; Sawin et al., 1992) and, thus, the phenotype can emerge due to assembly defects rather than post-mitotic nuclear separation. Because depletion of Klp61F led to a milder phenotype despite high knockdown efficiency, and because of the distinct localization of Feo and Klp3A in the aster-aster interaction zone (Fig. 1F,G), we followed up on the role of the latter two genes in maintaining internuclear distance

**Table 1:**
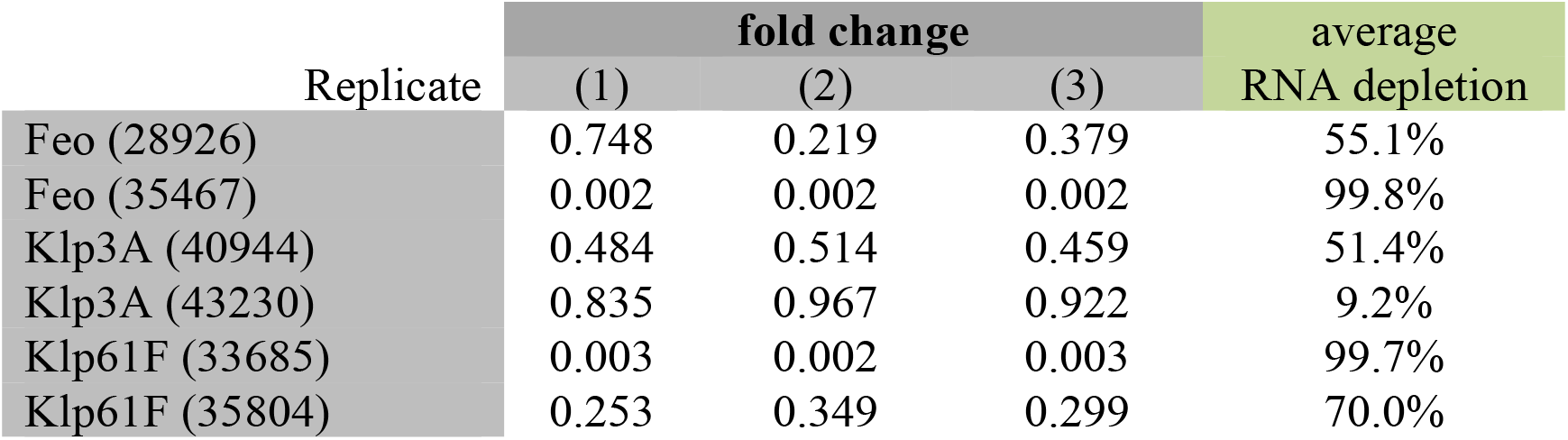
qPCR results of the RNAi expressing fly lines examined in this study.

| Replicate | fold change |  |  | average<br>RNA depletion |
| --- | --- | --- | --- | --- |
|  | (1) | (2) | (3) |  |
| Feo (28926) | 0.748 | 0.219 | 0.379 | 55.1% |
| Feo (35467) | 0.002 | 0.002 | 0.002 | 99.8% |
| Klp3A (40944) | 0.484 | 0.514 | 0.459 | 51.4% |
| Klp3A (43230) | 0.835 | 0.967 | 0.922 | 9.2% |
| Klp61F (33685) | 0.003 | 0.002 | 0.003 | 99.7% |
| Klp61F (35804) | 0.253 | 0.349 | 0.299 | 70.0% |

**Figure 2:**
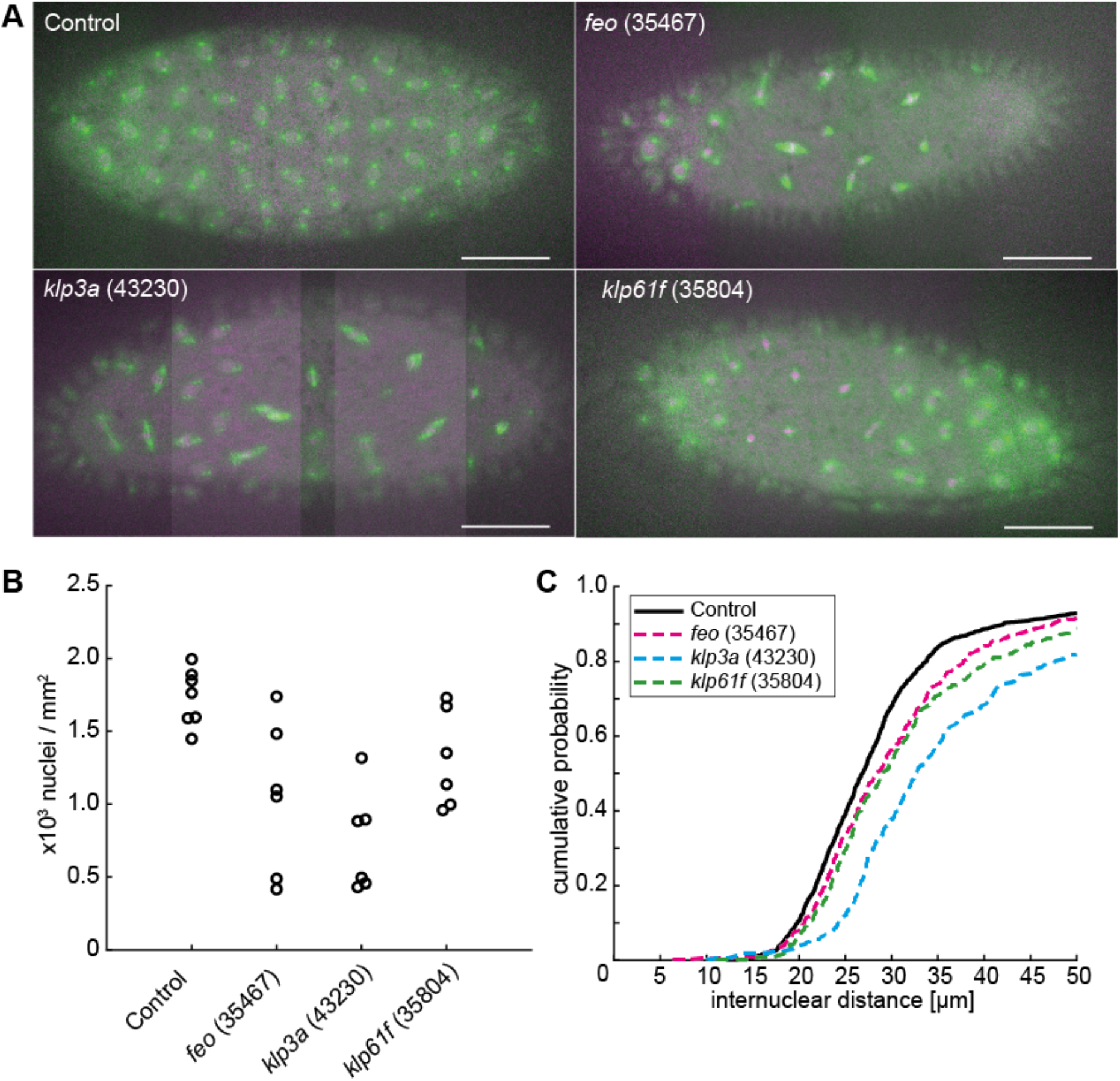
Knockdown of *feo, klp3a* or *klp61f* by RNAi leads to defective nuclear delivery to the embryo cortex. **A)** Maximum intensity projections from three-dimensional time-lapse movies of embryos expressing RNAi against *mCherry, feo, klp3a* or *klp61f*, and expressing Jupiter::GFP (green) marking microtubules and H2Av::RFP (magenta) marking chromatin. Knockdown embryos showed irregular nuclear distribution during the first interphase occurring at the cortex as compared to the regular nuclear distribution in control embryos (mCherry). Scale bar, 50 μm. **B)** A quantification of the number of nuclei per square millimeter showed a higher degree of variation between the six embryos knocked down for either of the three genes *feo* (35467), *klp3a* (43230) or *klp61f* (35804) as compared with control embryos. In all cases, the density decreased on average. Each data point represents one embryo. **C)** The cumulative probability function of the internuclear distance between first-order neighbors in embryos inhibited for Feo, Klp3A or Klp61F expression showed on average a higher internuclear distance. Thus, the number of nuclei at the cortex was smaller with broader distribution indicating greater irregularity and larger unoccupied areas with respect to the control. N=7 (control), N=6 (RNAi lines). Refer to Table 1, Suppl. Fig. 2 and Video 5.

### Developmental reset ex vivo reveals failure in nuclear distribution upon RNAi knockdown

Our analysis of nuclear distribution during knockdown in the embryo suggests that Feo and Klp3A are involved in nuclear delivery to the cortex. However, our knockdown approach *in vivo* has two drawbacks that can lead to misinterpretation: (i) The three proteins play a role in spindle midzone function, and their depletion may affect chromosome segregation in anaphase; (ii) The RNAi expression occurs chronically during late oogenesis. Thus, the irregular distribution of nuclei during cortical migration can be due to early sister chromatid separation errors, leading to missing nuclei in the embryo center and exponentially fewer in subsequent division cycles. Alternatively, inefficient nuclear separation following fertilization can lead to spindle fusion and mitotic errors. To circumvent the inability to detect accumulated effects, we performed time-lapse imaging of nuclear division cycles in embryo explants from preblastoderm embryos that were inhibited for either Feo or Klp3A protein expression. Because these explants contained only few dividing nuclei, we followed their distribution, or the failure thereof, while mimicking the very beginning of preblastoderm embryo development. We tracked individual nuclei undergoing division cycles and registered the distribution and any fusion events between sister and non-sister nuclei (Fig. 3A, Suppl. Video 6). Importantly, the time-lapse observation of nuclear divisions allowed us to determine if nuclear distribution changes arise from reduced separation or form mitotic failures and arrest, which has different consequences for the delivery of nuclei to the embryo cortex. In explants from control embryos, nuclei divide and distribute regularly in the entire explant (Fig. 3A, left, white dashed circle) until a saturated nuclear distribution is reached and occasional mitotic failures in the subsequent cycle are observed. The nuclear density at saturation is comparable to nuclear cycle 10 in the intact embryo (1800–2000 nuclei/mm^2^) (Foe and Alberts, 1983), corresponding to an internuclear distance of ∼25 μm (hexagonal approximation). Strikingly, the nuclei from *feo* and *klp3a* knockdown embryos also divide consecutively. The average distance between sister nuclei and between non-sister nuclei was lower in the test RNAi as compared to the control (Fig. 3B,C). However, the nuclear position after mitotic separation was maintained in the *feo* RNAi while knockdown of *klp3a* led to frequent spindle fusion at a comparable nuclear density and accumulation of mitotic failure. Interestingly, spindle length decreased upon depletion of *feo* (Fig. 3A, middle), but we did not observe significant decrease in spindle length upon *klp3a* depletion as reported earlier, most likely due to inefficient knockdown as compared to deletion (Williams et al., 1995). In summary, the inhibition of Feo expression leads to reduced nuclear separation between sister nuclei and incomplete occupation of nuclei within the explant, a hallmark of the unoccupied spaces in the blastoderm embryo (Fig. 2). However, while a knockdown of *feo* sustains mitotic divisions, *klp3a* knockdown produces a spindle fusion phenotype. It is conceivable that the reduction of Klp3A protein causes microtubule overlap over-growth and, despite crosslinking by Feo and other MAPs, these long overlaps allow for a net attractive force to build up between spindles (Gatlin et al., 2009). However, we cannot exclude that the partial knockdown of *klp3a* already causes mitotic defects related to spindle assembly and chromosome alignment. Because of these confounding effects we decided to focus on Feo exclusively.

**Figure 3:**
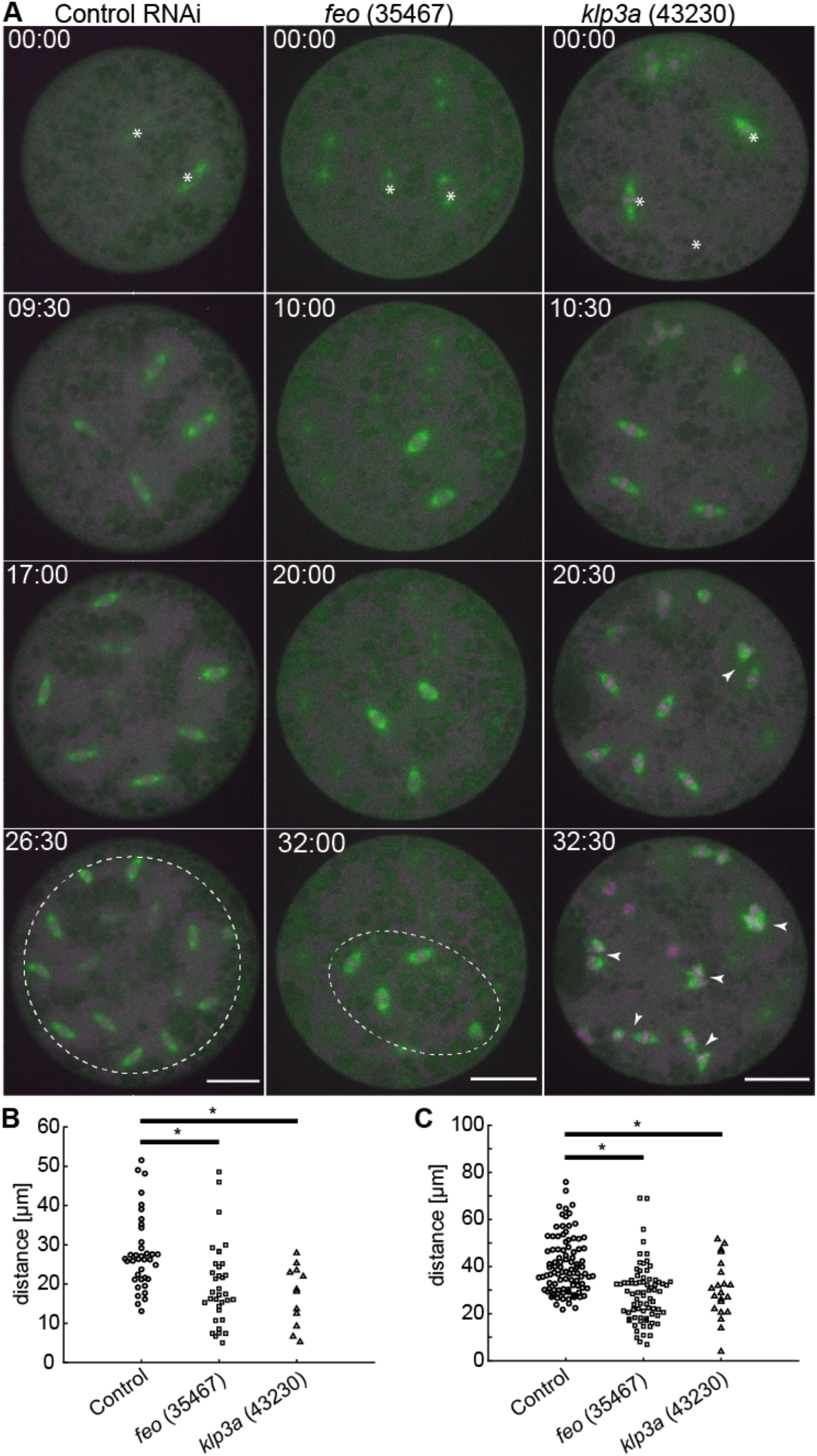
Knockdown of *feo, klp3a* or *klp61f* by RNAi leads to defective nuclear distribution in preblastoderm embryo explants. **A)** Maximum intensity projections from time-lapse movies of embryo explants under control conditions (*mcherry*) and RNAi against *feo, klp3a* or *klp61f* while expressing Jupiter::GFP (green) marking microtubules and H2Av::RFP (magenta) marking chromatin. Each panel shows metaphase of consecutive division. White stars in the first frame mark the position of dividing nuclei (sometimes out of focus). The control explants underwent normal nuclear divisions and distributed the daughter nuclei within the entire explant volume (dashed circle). Explants from *feo* RNAi embryos underwent mitotic nuclear divisions but daughter nuclei separate less efficiently, leading to a partial occupation of the cytoplasm (dashed ellipse). Explants from *klp3a* RNAi embryos underwent mitotic nuclear divisions with slightly less efficient distribution than in controls and with higher prevalence for spindle fusion (arrowheads). Scale bar, 30 μm; time in min:sec. Refer to Suppl. Video 6. **B)** Separation distance between daughter nuclei after mitotic nuclear division under control conditions and under knockdown for *feo* (35467) and *klp3a* (43230) in embryo explants. Separation distance was significantly reduced in both knock-down conditions (Control: N=4, n=38; *feo* (35467): N=2, n=36; *klp3a* (43230): N=3, n=23; p<0.01, Wilcoxon signed-rank test). **C)** Separation distance between first-neighbor non-sibling nuclei measured between mitotic divisions under control conditions and under knockdown for *feo* (35467) and *klp3a* (43230) in embryo explants. The separation distance was significantly shorter in both knockdown conditions (Control: N=3, n=98; *feo* (35467): N=3, n=77; *klp3a* (43230): N=3, n=50; p<0.05, Wilcoxon signed-rank test) though the effect is stronger under *feo* RNAi conditions.

### Displacement of nuclei is rescued in control but not in feo RNAi embryo explants

To test the model of an astral microtubule crosslinker-based separation mechanism for non-sister nuclei, we took advantage of the amenability of embryo explants for mechanical manipulation and designed an acute perturbation approach. We asked how Feo relocalizes when the distance between two interphase nuclei is manually reduced. Finally, we asked whether, under a *feo* knockdown condition, nuclei could adjust their position when brought in close proximity prior to division. To address these questions, we performed contact micromanipulation and changed the positions of two non-sister nuclei that were exiting mitosis (Fig. 4A, magenta, Suppl. Video 7), during the non-mitotic phase which is amenable to manipulation (Telley et al., 2012). Following manipulation of two nuclei (Fig. 4B, yellow) nuclei were typically in interphase, when Feo is not expected to localize due to cycle regulation. Nonetheless, we registered strong localization of Feo::mCherry in anaphase and telophase of the next cycle at midzones of all spindles, and strongly in the region between the manipulated nuclei (Fig. 4B bottom, starred) while asters from distant, non-manipulated nuclei did not recruit the microtubule crosslinker detectably (Fig. 4B, arrowhead). Next, we quantified nuclear separation of two neighboring nuclei dividing into four daughter nuclei by determining the four final positions (Fig. 4C), arranging these positions as a quadrilateral, aligning, annotating and overlaying them in a common coordinate system (Fig. 4D,E) and calculating area (Fig. 4F) and lateral distances (Fig. 4G,H). We performed these measurements under the control RNAi condition for nuclei in a large unoccupied cytoplasmic space, in a saturated space where several nuclei have spread through the entire explant, and in a crowded explant representing one more division cycle after saturation; this differentiated comparison takes into account nuclear density dynamics in cycling explants, as described in the previous section (Fig. 3). We found that the area of nuclear separation after manipulation is lower than in the non-manipulated and saturated space but indifferent from the crowded control (Fig. 4F). The manipulated nuclei divided and separated their daughter nuclei at ∼15 μm while the distance between non-siblings was maintained at ∼25 μm (Fig. 4G,H) (Telley et al., 2012), phenocopying the minimal separation seen in crowded explants for which distance maintenance is challenging. Interestingly, these separation distances are similar to what was reported for the blastoderm embryo (Kanesaki et al., 2011). Finally, we performed the manipulation of nuclear position in *feo* knockdown explants expressing Jupiter::GFP and H2Av::RFP. In these experiments, after manipulation, the daughter nuclei moved towards each other rather than apart, forming nuclear aggregates (Suppl. Fig. 3) phenocopying the treatment with Nocodazole as presented earlier (Suppl. Video 4). The separation of siblings was approximately the nuclear diameter (∼7 μm) (Fig. 4G, dashed line) and the separation of non-siblings was ∼10 μm (Fig. 4H). We conclude that acute repositioning of nuclei is detected by the separation machinery, as reported by Feo, and is counteracted to prevent spindle fusion or aggregation of nuclei. In other words, Feo is required to prevent nuclear collisions.

**Figure 4:**
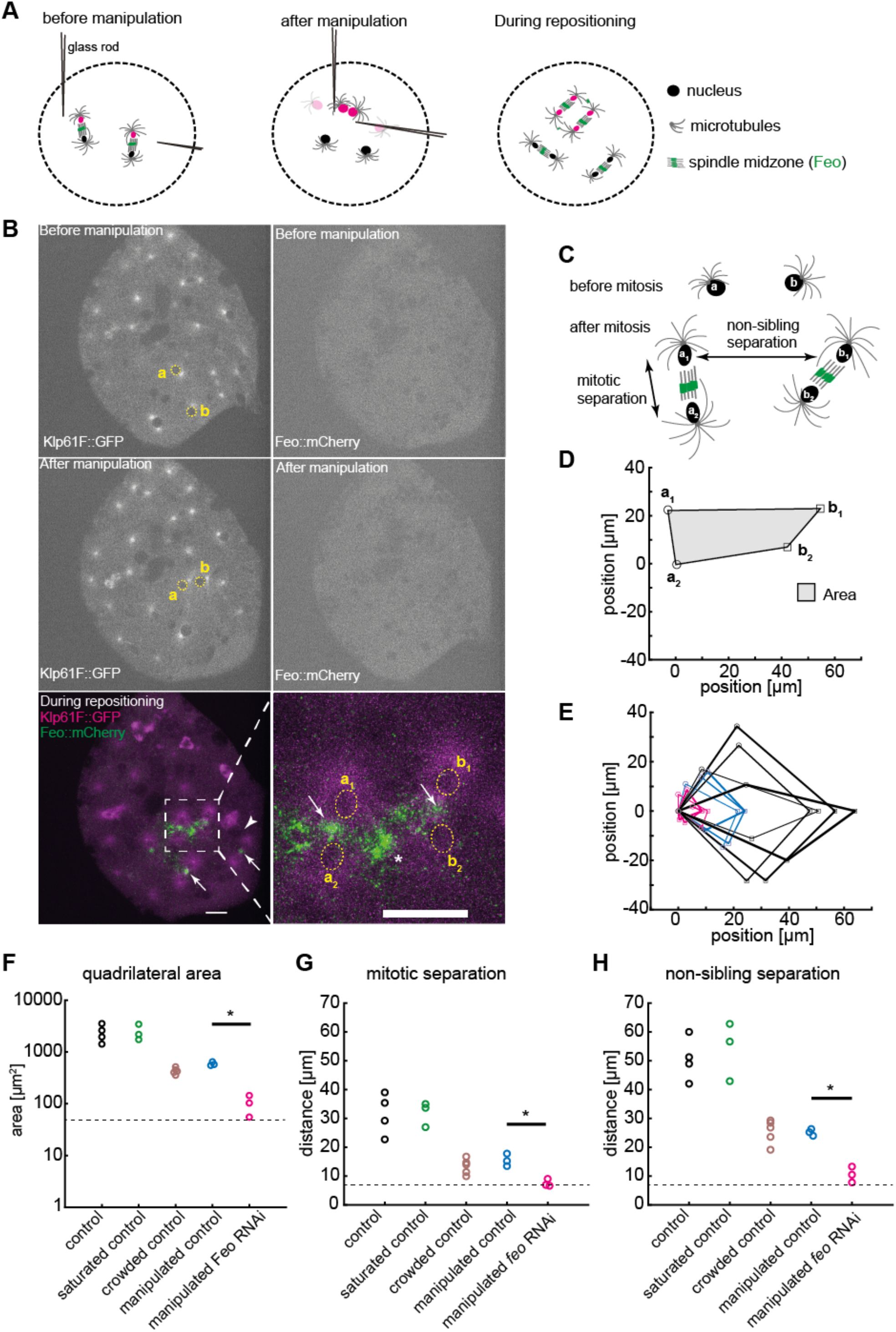
Explants fail to maintain nuclear separation distance following acute physical manipulation under *feo* knockdown. **A)** Scheme showing the manipulation of internuclear distance in embryo explants. After a mitotic division and nuclear separation, two non-sister nuclei (magenta) were brought close to each other during anaphase B–telophase by means of two glass rods. Subsequently, nuclei divided again, and daughter nuclei separated at defined distances. **B)** Fluorescence images illustrating physical manipulation of nuclear position in an explant made from an embryo expressing Klp61F::GFP (green) marking microtubules positively and nuclei negatively due to exclusion (dark disks), together with Feo::mCherry (magenta). The top row shows the GFP and mCherry signal before manipulation, the second row after manipulation. Physical manipulation decreased the distance selectively between two nuclei (labeled *a* and *b* and marked with yellow circles). In the subsequent mitosis and during repositioning of the daughter nuclei (bottom row), Feo localized at spindle midzones (arrows) and between the daughters of the manipulated nucleus (star in zoomed view) indicating that microtubule overlaps have formed. In contrast, Feo localization is not detectable between manipulated nucleus before or immediately after manipulation or between nuclei that have not been moved and are further apart (bottom, arrowhead). Scale bars, 15 μm. Refer to Suppl. Video 7. **C)** Schematic of the mitotic separation distance and non-sister separation distance. Nuclei a and b were brought close to each other and following a division give rise to daughters a_1_, a_2_, and b_1_, b_2_, respectively. **D)** Schematic of the quadrilateral area defined by the four nuclei a_1_, a_2_, b_1_, b_2_ after mitosis as shown in D). **E)** Overlay of quadrilaterals aligned for coordinate **a**_**2**_ and rotated so that the vector **b**_**1**_–**a**_**2**_ matches the *x*-axis. Control RNAi experiments without manipulation and with ample space in the explant are in black (N=4), experiments involving manipulation under control RNAi conditions are shown in blue (N=3), and manipulations experiments under knockdown of Feo are shown in magenta (N=3). The quadrilateral area for nuclei manipulated in *feo* RNA depleted explants was smaller when compared with control. **F)** Quadrilateral area for five different experimental conditions. The color code of panel E applies; additional control conditions without manipulation in explants almost saturated with nuclei (N=3) and in explants crowded with nuclei (N=5) are shown in green and brown, respectively. The dashed line designates the lower boundary where the four nuclei touch each other. The quadrilateral area for nuclei manipulated in *feo* RNA depleted explants was smaller when compared with control. **G)** The average mitotic separation distance between the dividing nuclei (|**a**_**1**_–**a**_**2**_|; |**b**_**1**_–**b**_**2**_|) was reduced in the manipulated *feo* RNAi condition and was close to the lower limit of separation (nuclear diameter) where the nuclei are touching each other. In contrast, sister nuclei were separated in all the control conditions. The color code is the same as in F. **H)** The average non-sister separation between the dividing nuclei (|**a**_**1**_–**b**_**1**_|; |**a**_**2**_–**b**_**2**_|) was reduced in the manipulated Feo RNAi condition and was close to the lower limit of separation where the nuclei are touching each other. In the control, the distance between the non-sister nuclei was ∼25 μm. The color code is the same as in F.

### Nuclear separation in the syncytium requires astral microtubule crosslinking by Feo

Feo is a dimer and, *in vitro*, has high affinity for binding two antiparallel microtubules (Bieling et al., 2010; Subramanian et al., 2013; 2010). In this function, Feo can generate a repulsive mechanical link – an apparent stiffness – which prevents concentric movement and eventual contact of neighboring nuclei. This model predicts a lower repulsion stiffness in the presence of a monomeric construct of Feo, which binds to the same microtubule lattice binding site as the full-length dimer but does not crosslink the antiparallel microtubules. We expect that this molecular competition leads to weaker crosslinking and can be measured as shorter internuclear distance, irregular separation or frequent nuclear contacts. Thus, we designed two protein expression constructs; one containing the full *feo* coding sequence (sFeoFL::GFP-His_6_), the other lacking the N-terminal, putative dimerization domain (Subramanian et al., 2013) (sFeoN::GFP-His_6_), and fused them to a C-terminal GFP and a His_6_ tag sequence (Fig. 5A,C). By design, the truncated construct should have unaltered microtubule-binding affinity while dimerization and, thus, cross-linking are abolished. Proteins were expressed in *E*.*coli*, affinity-purified and dialyzed into embryo explant compatible buffer (Telley et al., 2013) (Suppl. Fig. 4A). Full-length Feo::GFP forms dimers, as was reported for vertebrate homologs (Bieling et al., 2010; Subramanian et al., 2013; 2010), while the truncated protein is dimerization deficient and forms monomers or weak dimers, as shown in a native-PAGE (Suppl. Fig. 4B). When the full-length protein was injected at nanomolar concentration into embryos, GFP signal localized at the central spindle (Fig. 5B, arrow). As in transgenic embryos (Fig. 1F,G, arrowheads), we also detected small foci of green fluorescence between neighboring nuclei (Fig. 5B, arrowheads), suggesting that the purified protein and the transgenic construct localize identically and under cell cycle control with fluorescence disappearing in interphase and reappearing during telophase of the following cycle (Suppl. Video 8). Though less strong, the truncated protein construct also localized at the spindle midzone, however not in the cycle following injection but in the subsequent division cycle (Fig. 5D, Suppl. Fig. 4C, Suppl. Video 9). The low intensity suggests that the truncated construct competes weakly with endogenous Feo, and the cycle dependence indicates that dimerization is regulated by cell cycle kinases. Injection of full-length Feo::GFP maintained regular nuclear delivery to the embryo cortex while injection of truncated FeoN::GFP caused unnatural spindle contacts and nuclear separation defects (Suppl. Fig. 4C middle, Suppl. Video 8). Furthermore, when the full-length protein was injected into *feo* RNAi embryos the defective nuclear distribution was rescued to a large extent (Suppl. Fig. 4D). Nuclei arrived at the embryo cortex more symmetrically between anterior and posterior ends (Fig. 5E), in a less skewed distribution (Fig. 5F) and with more uniform internuclear distance (Fig. 5G) as compared to mock-injected *feo* RNAi embryos. Owing to the variability of injection and knockdown efficiency, we fully recovered nuclear density to a normal level in two embryos and significantly increased nuclear density in the remaining five embryos (Fig. 5H). Notably, the injected protein pool is stable for at least 90 minutes, throughout several division cycles. In summary, we show that a GFP-tagged full-length protein construct localizes correctly and rescues the gene knockdown in the germline. We conclude that it is functionally identical to the endogenous protein that is maternally deposited in the egg and stable during syncytial development.

**Figure 5:**
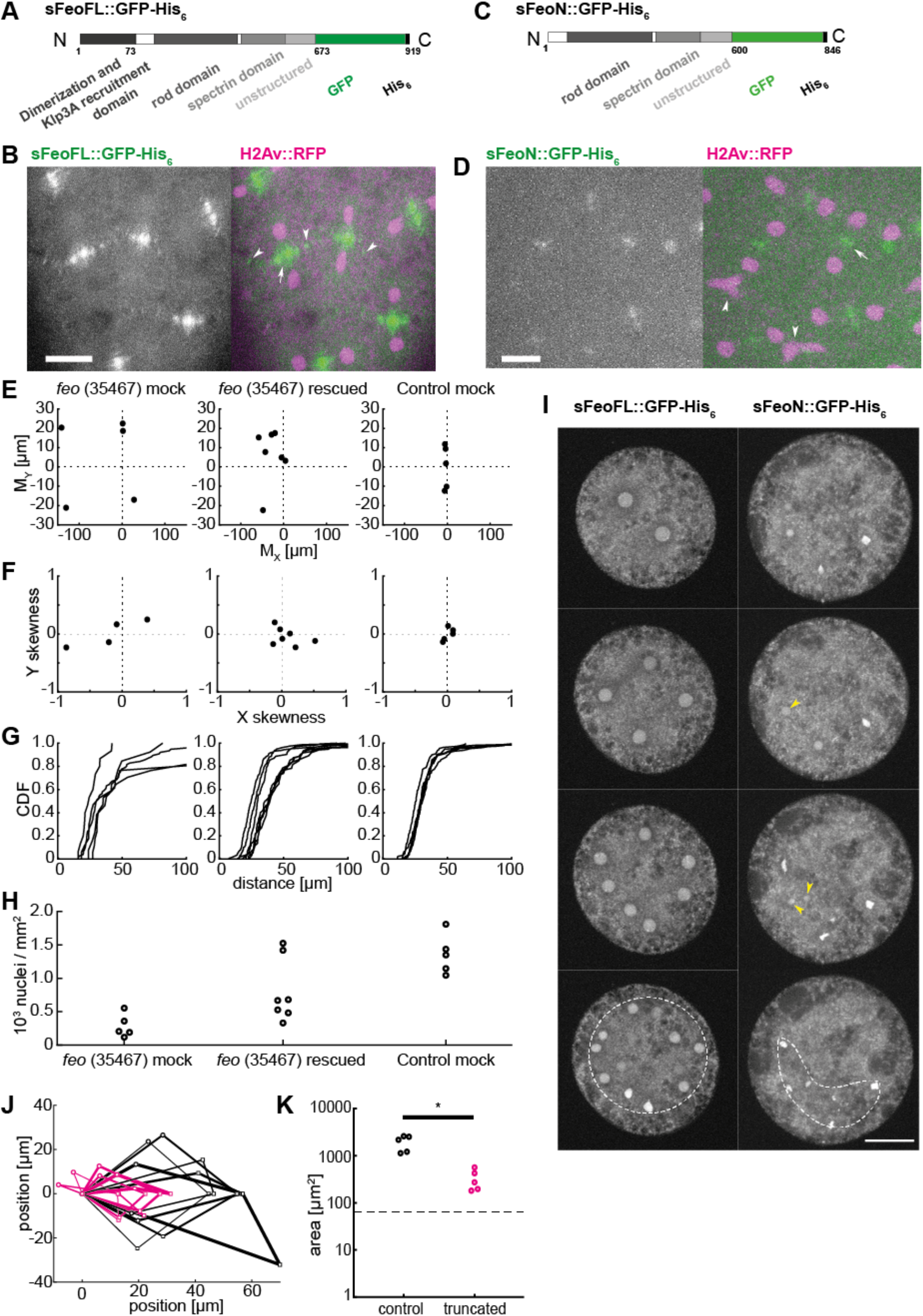
Purified Feo protein rescues nuclear separation in *feo* RNAi embryos, and a N-terminally truncated Feo abolishes nuclear separation. **A)** Scheme of the synthesized full-length Feo protein fusion construct containing a C-terminal GFP. The domains were determined based on sequence similarity from reported domains of the human construct. The N-terminal end induces dimerization and binds Klp3A, and the spectrin domain binds to microtubule lattice. **B)** Fluorescence image of the GFP-tagged full-length Feo protein in a (control) blastoderm embryo in telophase after protein injection. The GFP signal alone (left) is shown merged with H2Av::RFP in magenta (right). sFeoFL::GFP-His_6_ localized correctly, under cell cycle control, at the spindle midzone (arrow) and between daughter nuclei (arrowhead) as observed in the transgenic overexpression fly line shown in Fig. 1. Scale bar, 10 μm. Refer to Suppl. Video 8. **C)** Scheme of a truncated Feo construct lacking the first 73 amino acids of the putative dimerization and Klp3A recruiting domain, fused to a C-terminal GFP. **D)** Fluorescence image of the GFP-tagged truncated Feo protein in a (control) blastoderm embryo after protein injection. The GFP signal alone (left) is shown merged with H2Av::RFP in magenta (right). A faint GFP signal localized at the spindle midzone (arrow) in the second division after injection (Suppl. Fig. 4C). Nuclear separation defects manifest as neighboring nuclei touching or fusing after division (arrowheads). Scale bar, 10 μm. Refer to Suppl. Video 9. **E)** Plot of the 2-dimensional centroid vector (*M*_*X*_,*M*_*Y*_) of all cortical nuclei relative to the embryo center for Feo RNAi embryos either mock injected (left; N=5) or injected with sFeoFL::GFP-His6 protein (middle; N=7), compared with mock injected control (mCherry) RNAi embryos (N=5). The *x*-axis designates the anterior-posterior axis and the *y*-axis is the dorso-ventral axis of the embryo. Deviations from zero mark an acentric delivery of nuclei to the cortex. Along the anterior-posterior axis the injection of Feo full-length protein in Feo RNAi embryos partially rescued centering (middle) while mock-injected Feo RNAi embryos had anatomically eccentric nuclei (left), whereas mock-injected control (mCherry) RNAi embryos exhibited strong centering. **F)** Skewness plot of the positional distribution of all nuclei along the anterior-posterior (*x*) and dorsoventral (*y*) axis for the same conditions as in E. The asymmetric distribution in mock-injected Feo RNAi embryos (left) is partially rescued by Feo protein injection (middle) while mock-injected control embryos show little asymmetry. **G)** Cumulative distribution plot of the first-order neighbor distance between nuclei, for the same conditions as in e) and f). The irregular internuclear distances in mock-injected Feo RNAi embryos (left) are rescued to a considerable extent after full-length protein injection (middle) while mock-injected control (mCherry) RNAi embryos exhibit uniform inter-nuclear distances (right). **H)** The low nuclear density arriving at the cortex in mock-injected Feo RNAi embryos is partially rescued when full-length Feo protein is injected in preblastoderm Feo RNAi embryos. **I)** Addition of full-length Feo::GFP protein to embryo explants supports normal nuclear division and regular distribution within the explant space (left, white circle) while addition of truncated Feo protein reduces nuclear separation (arrowheads) causing occasional spindle fusion, and abolishes nuclear distribution (dashed envelope). Scale bar, 30 μm. **J)** Overlay of aligned quadrilaterals describing the nuclear separation after division in explants, as described in Fig. 4. Explants were generated from wildtype embryos and had ample space for the first few divisions. Experiments involving addition of full-length GFP-tagged Feo protein to the explant are in black (N=5), experiments involving addition of N-terminally truncated, GFP-tagged Feo protein are shown in magenta (N=5). **K)** The truncated Feo protein significantly reduced nuclear separation, as measured by the area of quadrilaterals shown in **J**, as compared to the full-length protein construct (black).

Finally, having designed and purified the truncated and the full-length protein with identical procedures, we asked how nuclear separation changes upon excess of the dimerization deficient Feo protein, added at 100–200 nM final concentration to wildtype embryo explants containing one or two nuclei. As control condition, we injected the full-length protein at the same final concentration into embryo explants, and despite this perturbation the explant supported normal nuclear separation and distribution (Fig. 5I, left). Conversely, adding the truncated protein construct worsened nuclear separation considerably after chromosomes segregated. Here, in contrast to the control condition, nuclei did not occupy the entire explant space after consecutive divisions. The short internuclear distance led to unnatural chromosome aggregation, fusion and eventually to mitotic failure. Nuclear separation of two neighboring non-sister nuclei, as measured by the quadrilateral area defined by their position, was significantly smaller than in control divisions in the presence of full-length Feo protein (Fig. 5J,K). We conclude that microtubule crosslinking by Feo generates a repulsive mechanical link between microtubule asters. Thus, it lies at the heart of nuclear separation maintenance during the multinucleated 1-cell stage of *Drosophila* embryo development.

## Discussion

A cornerstone of embryonic development is the formation of a polarized epithelium. Plants and many invertebrates achieve this developmental stage with a unicellular embryo undergoing nuclear proliferation followed by cellularization, a specialized form of cytokinesis (Hehenberger et al., 2012; Lecuit and Wieschaus, 2000). Recently, the molecular building blocks and morphogenetic characteristics of cellularization have also been identified as part of the life cycle of a non-animal eukaryote (Dudin et al., 2019). The offspring of *Sphaeroforma arctica* arises from nuclear proliferation, compartmentalization, and plasma membrane invagination generating a proto-epithelium from which newborn cells detach. These observations support the hypothesis that epithelia evolutionary predate animals (Dickinson et al., 2012). We propose that correct compartmentalization and generation of uninuclear offspring necessitates robust nuclear separation. If warranted true, then a separation mechanism must have coevolved with the origin of epithelia and was essential for the emergence of multicellularity.

Nuclear proliferation in a coenocyte poses a new challenge: How does the cell safeguard the separation and prevent contact of nuclei while their number increases? Two solutions seem plausible. On one hand, the cell may control the division axes and separate daughter nuclei along linear paths which do not cross. On the other hand, the cell may constrain internuclear distance independent of separation trajectories. Consider two nuclei that are about to divide and separate their progeny along the spindle axis (Fig. 6A). In a 3-dimensional space, none of the daughter nuclei may collide unless the spindle axes are both coplanar and non-parallel. Typically, nuclei migrate only 10–15μm away from the original spindle center before dividing again (Telley et al., 2012). This geometric constraint reduces configurations that produce colliding trajectories in a 2-dimensional topology to about 40% of all possible spindle axis orientations, so that axes intersect at an angle between zero (collinear) and 70º (Fig. 6B). Adding complexity, spindles in a network with optimal packing face a number of neighbors (6 in 2D, 12 in 3D) (Fig. 6C). Thus, a synchronously dividing spindle network will inevitably produce colliding trajectories of daughter nuclei. It is therefore necessary that, instead of controlling division axes, the cell controls nuclear proximity independent of the relative orientations they divide (Fig. 6B). This enables the syncytial embryo to divide hundreds of nuclei synchronously and distribute them to any unoccupied position. Here, we demonstrate a molecular mechanism that responds to short internuclear distances in the syncytium with a microtubule dependent repulsion. Each nucleus is associated with a radial array of microtubules nucleated by the centrosome, which duplicates and forms the two spindle poles in the next division. Prior, however, this microtubule aster guides nuclear migration and grows large enough to encounter microtubules from neighboring asters that migrate as well. This encounter leads to interdigitation of the microtubule plus–ends (antiparallel overlaps) and forms binding sites for crosslinking proteins. Our data shows that Feo, the Prc1 homolog in *Drosophila* and antiparallel microtubule crosslinker, plays a central role in defining a minimal internuclear distance in the syncytial *Drosophila* preblastoderm embryo.

**Figure 6:**
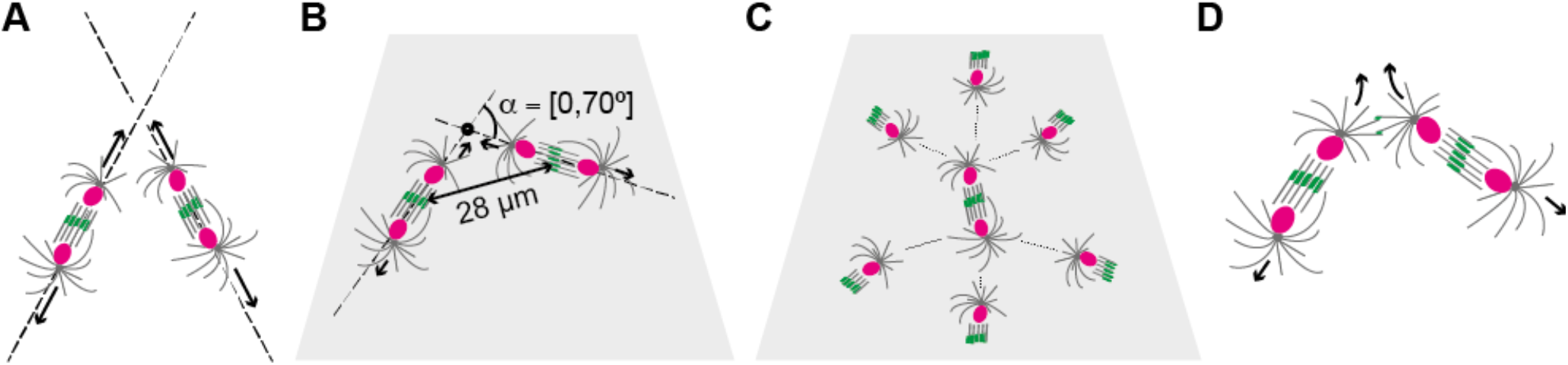
Feo and Klp3A prevent collision trajectories of dividing nuclei in space and on 2-dimensional topologies. **A)** Two neighboring spindles with division axes that are oblique. Nuclei separate along the spindle axis, which do not have an intersecting point and do not cause nuclear collision. **B)** Two neighboring spindles with coplanar spindle axes. If these axes are not parallel, they will always form an intersection point. However, because of the short nuclear migration from the previous spindle center (∼14 μm), the nuclear diameter (∼5 μm) and the average inter-spindle distance (∼28 μm), two non-sibling nuclei will only collide if the relative angle σ between spindle axes is ≤70º. **C)** In a two-dimensional topology of spindles with optimal packing each spindle has six neighbors. In this configuration, and considering the geometric constraints shown in **B**, any orientation of spindle axis for the spindle in the center will lead to collisions with non-sibling nuclei. **D)** Model of aster mediated repulsion between neighboring nuclei on a colliding trajectory after mitosis. Astral microtubule crosslinking by Feo and Klp3A generates a repulsive mechanical element that deviates the direction of separating nuclei from the spindle axis.

Vertebrate Prc1 is a microtubule binding protein with high turnover kinetics and at least 28 times higher affinity for antiparallel microtubule overlaps than for single microtubules (Bieling et al., 2010). This biochemical property, together with fluorescent labeling, renders Prc1 homologs reliable reporters for microtubule aster overlaps in live cell imaging assays (Nguyen et al., 2014). Prc1 crosslinking antiparallel microtubules generates a high affinity binding site for the motor protein Kinesin-4 (Kif4/Xklp1/Klp3A) at the overlap (Bieling et al., 2010; Kurasawa et al., 2004). In *Xenopus laevis* eggs and early embryos, where asters are unusually large, Prc1 and Kif4A define the radial organization and dynamics of microtubules and prevent invasion of neighboring asters by antiparallel microtubule crosslinking (Nguyen et al., 2018). The same protein module is responsible for recruiting cytokinesis signaling complexes and formation of the cleavage furrow (Nguyen et al., 2014). *In vitro*, in addition to maintaining a stable overlap length, co-activity of Prc1 and Xklp1 cause buckling of overlapping microtubules, which are immobilized at their minus end (Bieling et al., 2010). In a sliding assay of taxol-stabilized microtubules in which microtubules in solution and glass-immobilized microtubules form pairs cross-linked by Prc1, the antiparallel pairs of microtubules are slid apart by Kif4 (Wijeratne and Subramanian, 2018). This is reminiscent of plus–end directed sliding of Kinesin-5 (Eg5) (Kapitein et al., 2005) and explains the requirement of Prc1 orthologs for spindle elongation in several species (Khmelinskii et al., 2009; Schuyler et al., 2003; Vukušić et al., 2021; Wang et al., 2015; Zhu et al., 2006). Indeed, plus–end overlapping microtubules have an apparent mechanical stiffness that is governed by molecular friction and motor activity (Forth et al., 2014; Wijeratne and Subramanian, 2018). An assembly of tens of such microtubule pairs generates sufficient mechanical resistance against compressive forces in the nanonewton range, enough to keep two spherical organelles of 5–8 μm diameter and attached to the microtubule minus–ends (MTOC) well separated (Lele et al., 2018). Thus, modular upscaling of a single pair into overlapping radial arrays illustrates how the sliding mechanism of a Feo and Klp3A decorated antiparallel microtubule pair produces a repulsion between two syncytial nuclei.

Feo::GFP expressed in the transgenic line, or supplemented as purified protein, exhibited focal fluorescence signals in the blastoderm embryo and in the explant from preblastoderm embryos. Here, we showed that the length of these signal foci is surprisingly short and uniform. According to *in vitro* data, and neglecting any regulation other than affinity and stabilization activity for the underlying microtubule overlap to maintain such a short length, the concentration of Kinesin-4 in the cytoplasm must be at least one magnitude in excess of Feo (Bieling et al., 2010). Moreover, knockdown of *feo* by RNAi abolished the signal of Klp3A::GFP below detection, thus considerably reducing the bound fraction of Klp3A at the central spindle. In the embryo, while confirming their already established localization at the spindle midzone (D’Avino et al., 2007; Kwon et al., 2004; Page and Hawley, 2005; Wang et al., 2015; Williams et al., 1995), we recorded Klp3A::GFP signal colocalizing with Feo::mCherry in areas between neighboring spindle asters. However, we could not clearly assess the localization of Klp3A in explants from preblastoderm embryos due to the low signal intensity. A single-copy tagged Klp3A construct expressed with the endogenous promoter failed to provide sufficient signal, and we decided to work with overexpression constructs (Sarov et al., 2016). This indicates that the microtubule overlap–bound fraction of endogenous Klp3A is comparatively small despite the molar excess in the cytosol as derived from overlap length. Together, these observations point at a protein interaction network localized at antiparallel microtubule overlaps that is sensitive to small changes of Feo. As Feo binds microtubule overlaps independently (Bieling et al., 2010), the phenotypes in intact embryos and in explants can arise due to disproportionate Klp3A perturbation downstream of Feo. In summary, our live-cell microscopy data from blastoderm embryos and preblastoderm embryo explants support a ‘central spindle model’ built from individual pairs of microtubules crosslinked and length-regulated by Feo and Klp3A (Bieling et al., 2010). More importantly, we show how overlapping microtubules in the aster–aster interaction zone (Nguyen et al., 2014) form midzone–analogous cytoskeletal assemblies that persist throughout blastoderm development. This is particularly intriguing given that, at the embryo cortex from cycle 10 onwards, actin-based pseudo-furrows are thought to form pre-cellular compartments that prevent nuclear contact (Karr and Alberts, 1986; D. R. Kellogg et al., 1988; Lecuit, 2004; Mavrakis et al., 2009b). In the early blastoderm cycles, however, this compartmentalization may not yet be efficient enough to safeguard nuclear separation, and astral microtubule crosslinking persists as dominant mechanism. This interpretation is further supported by an earlier observation in mutants of the maternal-effect gene *sponge*, embryos of which do not form actin caps and pseudo-furrows in blastoderm stage but depict a homogenous nuclear distribution in cycle 10–11 (Postner et al., 1992).

Feo is essential for central spindle assembly and cytokinesis in somatic cells, containing two Cdk phosphorylation sites (Vernì et al., 2004). Feo, like Prc1 in human cells and Ase1p in fission yeast, is under cell cycle control and undergoes phosphorylation dependent localization from low intensity decoration of metaphase spindle microtubules to a strong localization at the central spindle in anaphase and telophase (Khmelinskii et al., 2009; Polak et al., 2017; Subramanian et al., 2013; Wang et al., 2015; Zhu et al., 2006). In the present work, we showed that the focal localization of Feo and Klp3A between neighboring nuclei is in synchrony with central spindle localization. It is in this phase of the division cycle when expanding spindles and separating nuclei cause a large spatial perturbation to the positional distribution (Kanesaki et al., 2011; Lv et al., 2020). Thus, a dual role for Feo under cell cycle control emerges; while it targets the central spindle at anaphase onset – forming the spindle midbody – it also binds to astral microtubule overlaps in a phase during which collision prevention is most needed.

In *Drosophila* embryos, spindle elongation at anaphase B is powered by the sliding activity of Klp61F (Brust-Mascher et al., 2009). Following the mechanism proposed by Baker et al. (1993), and because Klp61F is a candidate crosslinker and slider of overlapping astral microtubules, we performed RNAi knockdown in the germline. Inhibition of Klp61F expression led to lower density and non-uniform delivery of nuclei to the embryo cortex, confirming its essential role during preblastoderm development. However, owing to the established role of Klp61F in spindle assembly, the RNAi phenotype can emerge because of multiple chromosome segregation failures that were undetectable in the preblastoderm embryo. Here, the embryo explant assay overcomes an experimental limitation and enables time-lapse image acquisition of uni- or binuclear explants undergoing multiple divisions. Consequently, we could confirm that *klp61f* knockdown led to more frequent division failures rather than shorter nuclear separation. Still, Klp61F and Feo can functionally cooperate in crosslinking astral microtubules because both proteins recognize and bind to microtubule pairs, though with different preference for microtubule orientation (Bieling et al., 2010; Kapitein et al., 2005; E. H. Kellogg et al., 2016; Subramanian et al., 2010). In human cells, Prc1-dependent Kif4A motor activity and the microtubule sliding by Eg5 are redundant for spindle elongation during anaphase (Vukušić et al., 2021). Interestingly, in *Drosophila*, while Feo modulates binding and localization of Klp61F at the spindle midzone in anaphase, Klp61F cannot functionally rescue the absence of Feo (Wang et al., 2015). Presumably, Ase1p/Prc1/Feo binding to microtubule overlaps creates a protein binding hub for motors and regulators (Bieling et al., 2010; D’Avino et al., 2007; Hu et al., 2012; Khmelinskii et al., 2009; Sasabe and Machida, 2006; Subramanian et al., 2013). This property has not been demonstrated for Kinesin-5 orthologs. Together, the collection of our and other evidence suggests that Klp61F is not at the core of astral microtubule driven nuclear separation.

Lastly, the reader may wonder how astral microtubule overlap crosslinking by Feo and Klp3A defines the internuclear distance metric, leading to a distribution of syncytial nuclei with high regularity. In an earlier study, Telley et al. (2012) showed that microtubule aster size varies throughout the nuclear division cycle, reaching a maximum of 11 ± 3 μm in telophase. Herein, the aster size represents the length distribution of microtubules which, for dynamic microtubules with non-growing minus–end, is well approximated with an exponential distribution (Howard, 2001). We assume that two microtubules from neighboring asters grow at least to average length, overlap with their plus–ends and are collinear. If the stabilized overlap length is ∼1 μm, then the total length from centrosome to centrosome is on average 21 ± 4 μm. Considering that a centrosome is ∼1 μm large, and that a nucleus in late telophase is 5 ± 1 μm in diameter, the total distance between the centers of neighboring nuclei is 28 ± 4 μm. This estimate is in good agreement with the internuclear distance distribution measured from center to center of each nucleus (Fig. 3C), the minimal non-sibling internuclear distance in extract (Fig. 4H) and earlier reported separation distances of daughter nuclei (Telley et al., 2012). Thus, the short antiparallel overlap length of microtubules from neighboring asters and the microtubule length distribution are sufficient to explain the geometry of nuclear distribution in the *Drosophila* syncytial embryo.

## Materials and Methods

### D. melanogaster

Rearing of flies for general maintenance was done as previously described (Stocker and Gallant, 2008). The following fly lines were used to make recombinants or trans-heterozygotes: Jupiter::GFP (BDSC# 6836), Jupiter::mCherry (generated by and obtained from Nick Lowe in D. St Johnston’s lab, The Gurdon Institute), pUbq>Spd2::GFP (gift from M. Bettencourt-Dias, IGC), H2Av::RFP (BDSC# 23650), Feo::GFP (BDSC# 59274), Feo::mCherry (BDSC# 59277), Klp61F::GFP (BDSC# 35509), Klp3A::GFP (VDSC# 318352), RNAi targeting *feo* (BDSC# 28926 and 35467), RNAi targeting *klp3a* (BDSC# 40944 and 43230), RNAi targeting *klp61f* (BDSC# 33685 and 35804), RNAi targeting *mcherry* (BDSC# 35785), UASp–GFP (BDSC# 35786).

### RNAi experiments

Knockdown experiments were performed using the TRiPGermline fly lines for RNAi in germline cells (Perkins et al., 2015). The UAS-hairpin against a gene of interest was expressed using V32– Gal4 (gift from M. Bettencourt Dias) at 25ºC. The expression profile of V32–Gal4 in the oocyte was assessed by dissecting ovaries of flies expressing UASp–GFP at 25ºC and comparing GFP expression at different developmental stages with fluorescence microscopy.

### Sample preparation and extraction

Embryos were collected from apple juice agar plates mounted on a fly cage. They were dechorionated in 7% sodium hypochlorite solution, aligned and immobilized on a clean coverslip using adhesive dissolved in heptane and covered with halocarbon oil (Voltalef 10S). Extraction of cytoplasm from individual embryos and generation of explants was performed on a custom-made microscope as previously described (de-Carvalho et al., 2018; Telley et al., 2013).

### Quantitative PCR

To measure the transcript levels of *feo, klp3a* and *klp61f*, total RNA was extracted following standard procedures (PureLink RNA Mini Kit, Ambion) from embryos collected after 40 minutes of egg laying. A cDNA library was made from Oligo(dT)12–18 as described in the manufacturer’s protocol (Transcriptor First Strand cDNA Synthesis Kit, Roche). Quantitative PCR was performed using Quantifast SYBR Green PCR Kit (204052) and QuantiTect Primers for *feo* (QT00919758) in *feo* RNAi (35467 and 28926), *klp3a* (QT00497154) in *klp3a* RNAi (40944 and 43230) and *klp61f* (QT00955822) in *klp61f* RNAi (35804 and 33685). *actin* (QT00498883) was used as a housekeeping gene control. Results are presented in Table 1.

### Purification of sFeoFL::GFP and sFeoN::GFP

The full coding sequence of the *feo* gene fused to a C-terminal GFP tag, was synthesized and codon optimized by NZYTech, referred to herein as sFeoFL::GFP. The DNA was cloned into the vector pET-21a containing a C-terminal His_6_-tag, and transformed into *E*.*coli Rosetta* cells. The coding sequence of the *feo* gene without the initial 73 N-terminal residues, referred here as truncated sFeoN::GFP construct, was amplified from the synthesized sFeoFL::GFP construct and re-cloned into the pET-21a vector. Both proteins were produced by IPTG induction at 25ºC. After 4h of incubation, the cells were harvested and resuspended in lysis buffer (100mM K-HEPES pH 7.4, 500 mM NaCl, 10% glycerol, 0.1% Triton X-100, 3 M urea, supplemented with protease inhibitors (Roche) and 100U of DNAse type I (NZYTech)). The cells were lysed using the digital sonifier® (SLPe, Branson) at 70% amplitude with 6 pulses of [30 sec on]–[30 sec off] and clarified by centrifugation at 30,000g for 45 minutes at 4ºC. For purification of the truncated construct, the supernatant was loaded onto a 5 ml HiTrap Chelating HP (GE Healthcare) charged with 0.1 mM NiCl_2_ and equilibrated with wash buffer (100 mM K-HEPES pH7.4, 500 mM NaCl, 10% glycerol, 40 mM imidazole, 1 mM 2-mercaptoethanol), extensively washed with this buffer and eluted with elution buffer (100 mM K-HEPES pH7.4, 500 mM NaCl, 10% glycerol, 500 mM imidazole, 1 mM 2-mercaptoethanol) throughout a gradient of 6 CV. For purification of the full-length construct, the supernatant was loaded onto a 1 ml HiTrap TALON crude (GE Healthcare) charged with 50 mM CoCl_2_ and equilibrated with wash buffer (100 mM K-HEPES pH 7.2, 500 mM NaCl, 10% glycerol, 5 mM imidazole, 1 mM 2-mercaptoethanol), extensively washed with this buffer and eluted with elution buffer (100 mM K-HEPES pH 7.2, 500 mM NaCl, 10% glycerol, 150 mM imidazole, 1 mM 2-mercaptoethanol), throughout a gradient of 20 CV. Fractions containing the protein of interest were pooled, the buffer exchanged into embryo explant compatible buffer (100 mM K-HEPES pH 7.8, 1 mM MgCl_2_, 100 mM KCl) using a PD-10 desalting column (GE Healthcare) and concentrated using a 50K MWCO Amicon^®^ Ultracentrifugal filter (Merck). The purifications were performed using the ÄKTApurifier protein purification system (GE Healthcare) and the chromatographic profile of both proteins was followed by measuring the absorbance at 280 nm, 254 nm and 488 nm in the UV-900 monitor. The size exclusion method resulted in Feo constructs strongly associated to an unknown contaminant at ∼50 kDa. The concentration of each construct was estimated ∼50% of the total measured protein concentration based on band analysis of SDS-PAGE. Total protein concentrations were measured with a NanoDrop2000 UV-Vis spectrophotometer (ThermoFisher).

### SDS PAGE, Native PAGE and Blotting with quantification

Polyacrylamide gels for electrophoresis and Coomassie-staining were made with 15% SDS. Molecular mass was estimated by linear regression of log_10_(M_W_ [Da]) as a function of the migration distances [cm] of the ladder (BioRad, 161-0375). To determine the dimerization of sFeoFL::GFP and sFeoN:GFP a Western Blot was generated from a 4-12% gradient SDS and a Native polyacrylamide gel side-by-side using a mouse anti-GFP antibody (Roche Nr. 11814460001) at 1:500 dilution. The molecular mass of the protein samples was estimated using a linear regression of log_10_(M_W_ [Da]) as a function of the migration distances [cm] of the ladder (NZYTech MB090); in the SDS gel a regression was performed on the entire ladder while for the Native gel only the higher molecular mass ladder (100–245 kDa) was considered.

### Addition of exogenous purified proteins

Purified porcine Tubulin (Cytoskeleton) was labeled with Alexa-647 (Invitrogen, ThermoFisher) following a published protocol (Hyman, 1991) and injected into embryos or explants at 0.3–0.8 mg/ml. Freshly purified sFeoFL::GFP and sFeoN::GFP were injected at 2 mg/ml in EC buffer in embryos or explants. This concentration was a result from a series of titrations over a magnitude of different concentrations and assessing phenotype or rescue. For embryos, the injected volume assumed a spherical shape with diameter D ≈ 0.018 mm, resulting an injection volume of 3.05 × 10^−6^ mm^3^. The average length and width of the embryo are 0.5 mm and 0.2 mm, respectively (Markow et al., 2008). Assuming an ellipsoid geometry for the embryo, its volume is ∼10^−2^ mm^3^. Thus, the final concentration of injected protein after equilibration in the entire embryo was 5–6 nM. For explants, both protein constructs were added to explant cytoplasm at 1:200 (vol/vol), resulting in a final concentration in the cytoplasm of 100–200 nM. Importantly, such an excess of full-length Feo protein preserved nuclear divisions and distribution.

### Treatment of embryo explants with Nocodazole

Nocodazole was dissolved in DMSO and diluted to 200 μM in EC buffer at pH 7.8, with a concentration of DMSO in EC buffer of 2 % (v/v). This buffer mixture was added to explants at ∼1:50 (v/v); this was estimated by measuring the diameters of the buffer droplet in the explant and the explant itself, calculating their area and scaling according to area change (Telley et al., 2013). The final concentration of Nocodazole in the explant was ∼4μM. Control experiments were conducted by adding the buffer mixture without the drug to explants at similar ratios.

### Image acquisition, processing and analysis

Transmission light microscopy images were obtained with a 10x 0.25NA objective, and the polarizer and analyzer of the microscope in crossed configuration. Time-lapse confocal fluorescence Z stacks were acquired on a Yokogawa CSU-W1 spinning disk confocal scanner with 488 nm, 561 nm and 640 nm laser lines. Images of whole embryos were acquired with a 40x 1.3NA oil immersion objective. Images of embryo explants were acquired with a 60x 1.2NA or a 40x 1.15NA water immersion objective. Images were recorded with an Andor iXon3 888 EMCCD 1024×1024 camera with 13 μm square pixel size, and a 2x magnification in front of the camera. Image processing i.e. making Z-projections, image cropping, image down-sampling, and video generation, was performed in Fiji (Schindelin et al., 2012). Whole embryo images for knockdown experiments were obtained by pairwise stitching using a plugin in Fiji.

The fluorescence signal of Feo::GFP in explants was analyzed with the line profile tool in Fiji. First, images of dividing nuclei during anaphase or telophase were filtered with a Gaussian kernel (σ = 1.2). Spot-like signals located between non-sibling nuclei were identified and, where spots were non-circular, a line was drawn along the longer axis. The angle of the line relative to the image coordinate system was recorded, and an intensity profile was generated. Profiles were aligned relative to the position of highest intensity and averaged. For each image, an intensity profile from a location void of microtubule signal was generated to obtain the background and the standard deviation of Feo::GFP intensity. Finally, the size of the spot was determined by calculating the width of the curve where the intensity was higher than two times the standard deviation of the background. The angle of every profile line was transformed relative to the closer of the two axes that connect the centrosome of one nucleus with the centrosome of the two neighboring sister nuclei (Fig. 1C) termed as angle *θ*. A probability density plot from all measured angles was generated in MATLAB^®^.

The Voronoi segmentation was performed by determining the position of the spindle poles manually in Fiji and providing these coordinates to the voronoi function implemented in MATLAB^®^.

The nuclear density in whole embryo images was obtained by measuring the area of the visible part of the embryo after manually tracing the border and dividing the number of nuclei by this area. The localization of nuclei in whole embryos was performed manually in Fiji. The precision of localization was 0.25 μm (intra-operator variability). Localization coordinates were imported into MATLAB^®^ and transformed with respect to the coordinate system of the embryo, as defined by the anterior pole as coordinate origin, and the anterior-posterior axis as *x*-axis. The first-order internuclear distances were obtained from the triangulation connectivity list (‘delaunay’ function), while excluding any edge connections, and by calculating the distance between the remaining connections. The cumulative distribution function of internuclear distances from individual embryos was obtained with the ‘ecdf’ function in MATLAB^®^. An average cumulative distribution function from several embryos was generated after pooling all distances together. Next, the deviation of the centroid of nuclear positions from the anatomical center of the embryo was obtained using the formula:

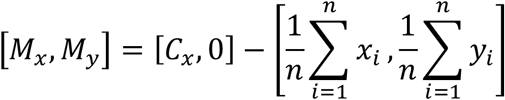

whereby an estimate for the anatomical center of the embryo, [σ*x*, 0] with respect to the embryo coordinate system, is given by half the pole-to-pole distance on the *x*-axis and zero on the *y*-axis. The third-order moment of the distribution of nuclear coordinates was calculated with the ‘skewness’ function in MATLAB^®^, providing a measure for left-right asymmetry.

The measurement of inter-nuclear distances in embryo explants was performed manually in Fiji. The precision of distance measurement was ±0.12 μm as determined by repeated measurement (intra-operator variability). The intensity profile plots of Klp3A::GFP in the Feo RNAi background were obtained using the line profile tool in Fiji, by drawing a line between daughter nuclei in the red (H2Av::RFP) channel and generating an intensity profile plot in the green channel, aligning these profiles according to the peak intensity and averaging profiles from different locations and embryos.

Plots of aligned quadrilaterals were generated with MATLAB^®^ by coordinate transformation. The area was calculated using Gauss’ trapezoidal formula for general polygons:

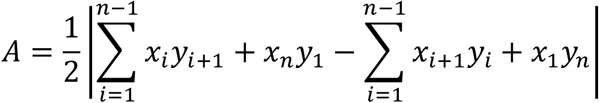

while *n* = 4 for ‘quadrilaterals’. For each quadrilateral, representing two sets of dividing nuclei, the average of the two involved mitotic separation distances and the average of the two involved non-sibling separations were calculated and plotted. All graphs were made with MATLAB^®^.

### Statistical analysis

We have plotted, unless otherwise mentioned, the distribution of raw data or cumulative probabilities. Sample size (n) and the number of experimental repeats (N) are reported in the figure legends. A Wilcoxon rank-sum test was performed with MATLAB^®^ starting with a significance level *α* = 0.05.

## Acknowledgements

We thank members of the Telley lab for fruitful discussions, Jonathon Scholey for constructive comments on the manuscript and for fly stocks, and Thomas Surrey for discussions throughout the project. We thank the staff of the Fly Facility, the Advanced Imaging Facility and the Technical Support Service at the Instituto Gulbenkian de Ciência (IGC). Transgenic fly stocks were obtained from the Vienna Drosophila Resource Center and Bloomington Drosophila Stock Center (NIH P40OD018537). We acknowledge financial support provided by Fundação Calouste Gulbenkian (FCG), European Commission FP7 Marie Curie CIG to I.A.T. (PCIG13-GA-2013-618743), Human Frontiers Science Program YIG to I.A.T. (RGY0083/2016), a doctoral fellowship SFRH/BD/52174/2013 to O.D. from Fundação para a Ciência e a Tecnologia (FCT). We acknowledge LISBOA-01-0145-FEDER-007654 supporting IGC’s core operation, LISBOA-01-0145-FEDER-022170 (*Congento*) supporting the Fly Facility, and PPBI-POCI-01-0145-FEDER-022122 supporting the Advanced Imaging Facility, all co-financed by FCT (Portugal) and Lisboa2020, under the PORTUGAL2020 agreement (European Regional Development Fund).

## Author contributions

OD, JC and IAT conceived and designed the project. OD and IAT designed experiments and OD performed them with occasional support from JC. DMV and OD designed, purified and characterized the protein constructs. IAT designed and assembled the instrument for explant generation and manipulation. OD, JC and IAT prepared the figures and wrote the manuscript.

## Supplementary Figures

Supplementary Figures 1 – 5

**Supplementary Figure 1.**
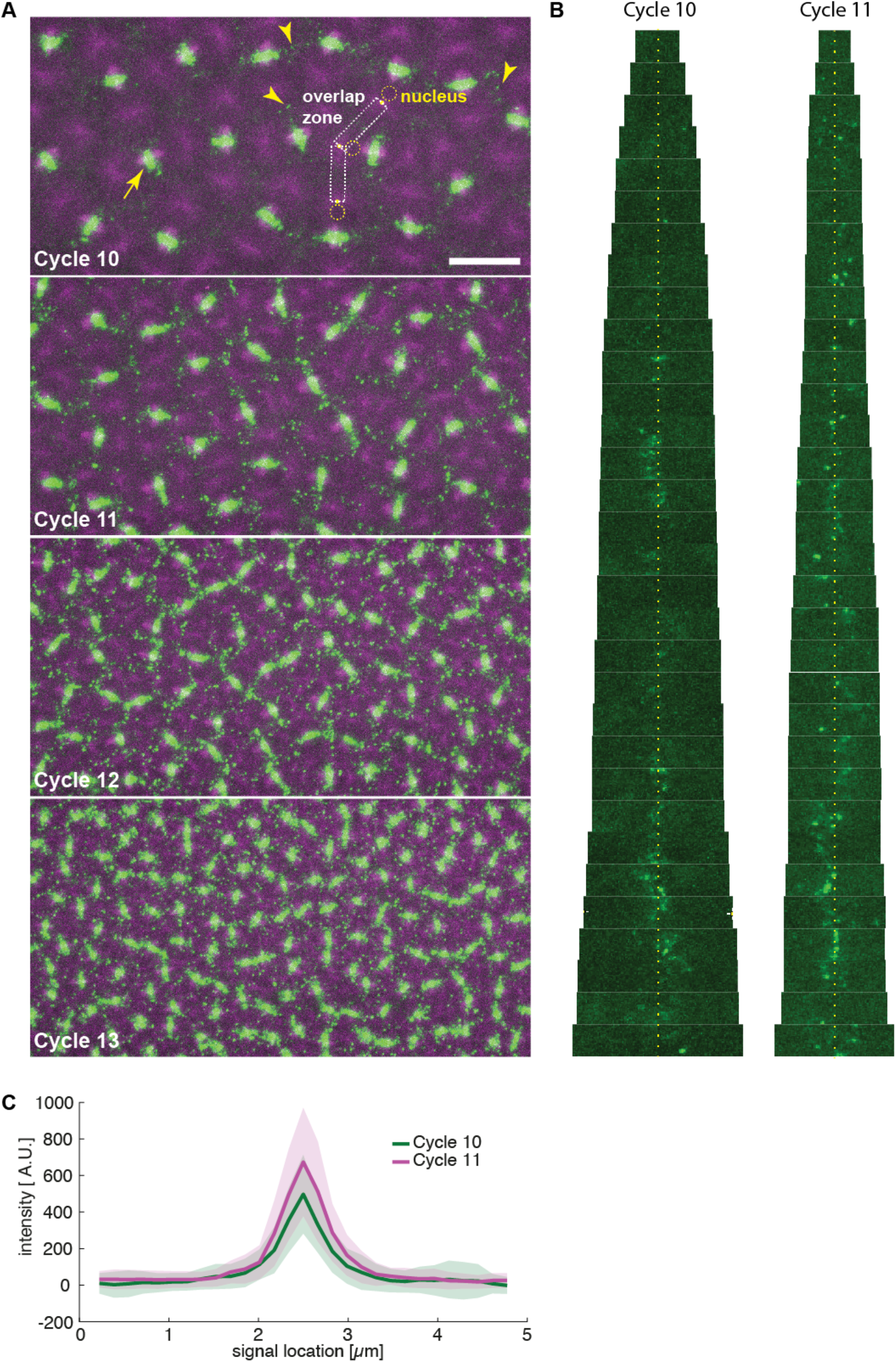
related to Figure 1: Feo localization between neighboring nuclei in the developing embryo. **A)** Two-color still images of a blastoderm embryo expressing RFP::β-tubulin (magenta) and Feo::GFP (green) during cycles 10,11,12 and 13, showing Feo localization between sister nuclei as part of the spindle midzone (arrow) and between neighboring non-sister nuclei as distinct foci (arrowheads). While in cycle 10 very few Feo foci are visible, their appearance and density increase with every further division cycle. Qualitatively, their size does not change. For further quantification, we tracked nuclei (dashed circle) – visible by the circular absence of tubulin signal – and the spindle pole (yellow dot) – the peak signal in the tubulin channel next to the nuclei – and cropped the image into putative microtubule overlap zones (dashed rectangles). Scale bar, 20 μm. **B)** Sequence of fluorescence images of Feo::GFP cropped from the putative overlap zones (see panel A), ordered by size i.e. distance between neighboring spindle poles, for cycles 10 and 11 (N=3 embryos). As expected, neighbor distance is overall smaller in cycle 11. Note that signals appear predominantly in the center of the overlap zones (yellow dotted line) representing overlaps equidistantly from each neighbor spindle pole. **C)** Quantification of foci size embryos of cycle 10 and 11 (where foci can still be reliably distinguished). The average size of foci is indistinguishable for these two cycles and matches the size of foci observed in explants (Fig. 1D). Shaded areas represent s.d. (N=39).

**Supplementary Figure 2.**
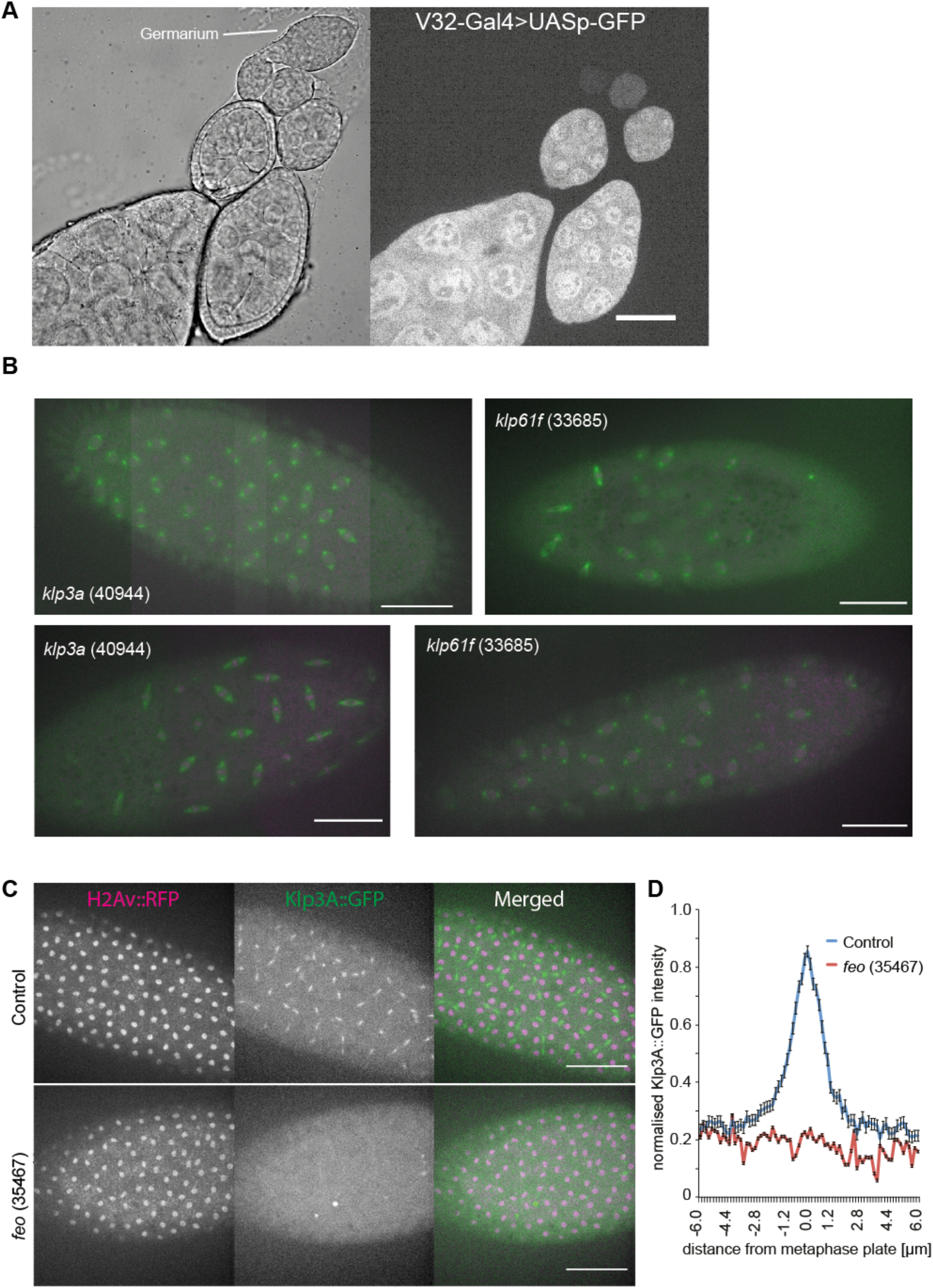
related to Figure 2: **A) V32-Gal4 drives expression during late oogenesis**. A construct expressing V32–Gal4 driving UASp–GFP expression specifically in the female germline indicates that the peak expression of GFP is achieved only at late stages of oogenesis. It illustrates the expression pattern of UASp constructs under the same Gal4 driver, including the various RNAi constructs described here have maximum effect in late oogenesis. Scale bar, 10 μm. **B) Knockdown of Klp3A (40944) or Klp61F (33685) by RNAi leads to defective nuclear delivery to the embryo cortex**. Maximum intensity projections from three-dimensional time-lapse movies of embryos inhibited for Klp3A or Klp61F expression by RNAi while expressing Jupiter::GFP (green) marking microtubules and H2Av::RFP (magenta) marking chromatin. These two alternative RNAi expressing fly lines exhibited similar irregularity in nuclear distribution during the first interphase occurring at the cortex. Scale bar, 50 μm. **C-D) Depletion of Feo abolishes recruitment of Klp3A to the spindle midzone. C)** Snapshots from a time-lapse of embryos expressing H2Av::RFP (magenta, left panel) and Klp3A::GFP (green, middle panel) during anaphase B or telophase. Feo knockdown embryos failed to recruit Klp3A at the spindle midzone when compared with the control embryos expressing no *feo* RNAi. **D)** Quantification of Klp3A::GFP intensity measured at the spindle midzone along the spindle axis in control and Feo RNAi (35467) embryos. Scale bar, 50 μm.

**Supplementary Figure 3.**
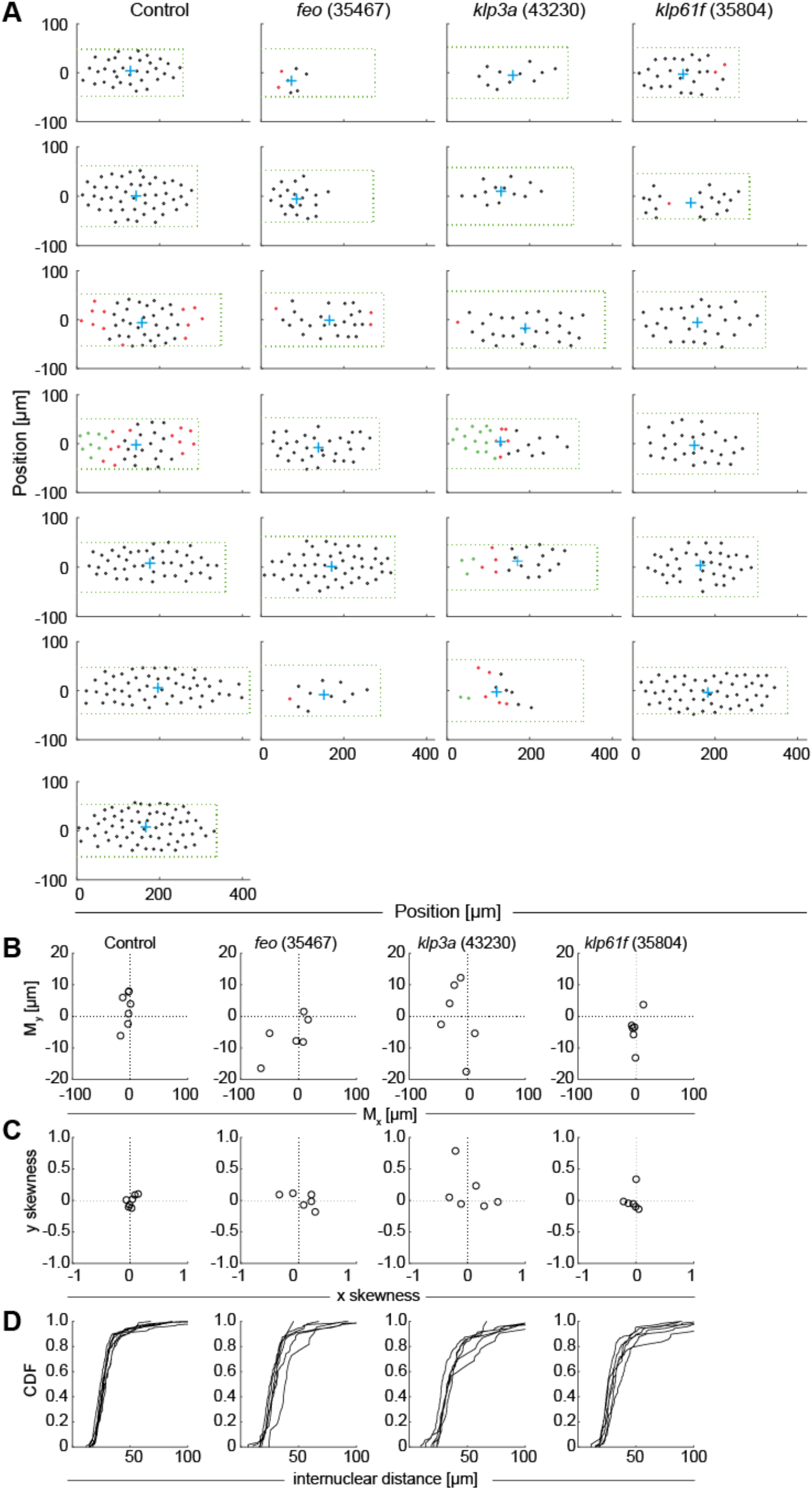
(related to Figure 2): Positions of nuclei in each of the analyzed embryos and distribution measurements highlight irregularity in the knockdown constructs. **A)** The position (circle) of every nucleus arriving at the embryo cortex after the last preblastoderm division, relative to the axial and lateral borders of the embryo, for each condition – Control (mCherry), Feo (35467), Klp3A (40320), Klp61F (35804). The green dashed rectangle represents the area of the embryo bounded by the length and width of the visible embryo in the confocal stacks, with the anterior end at the coordinate origin. The blue cross represents the location of the 2-dimensional centroid determined from the position of all nuclei. The nuclei in interphase of the first division at the cortex are marked in black, the nuclei that have progressed to metaphase / anaphase are marked in magenta, and the nuclei in telophase / (next) interphase are marked in green. Note that the RNAi expression and penetration exhibit a variability, which we attribute being responsible for the range of phenotypes. **B)** Plot of the 2-dimensional centroid vector (*M*_*x*_,*M*_*y*_) of all cortical nuclei relative to the embryo center. The *x*-axis designates the anterior-posterior axis and the *y*-axis is the dorsoventral axis of the embryo. Deviations from zero mark an acentric delivery of nuclei to the cortex. **C)** Skewness plot of the positional distribution of all nuclei along the anterior-posterior (*x*) and dorsoventral (*y*) axis. Feo RNAi and Klp3A RNAi embryos showed asymmetric nuclear distribution while nuclei in Klp61F RNAi embryos were distributed symmetrically. **D)** Cumulative distribution plot of the first-order neighbor distance between nuclei. All RNAi lines showed higher variability in internuclear distance as compared with the control.

**Supplementary Figure 4.**
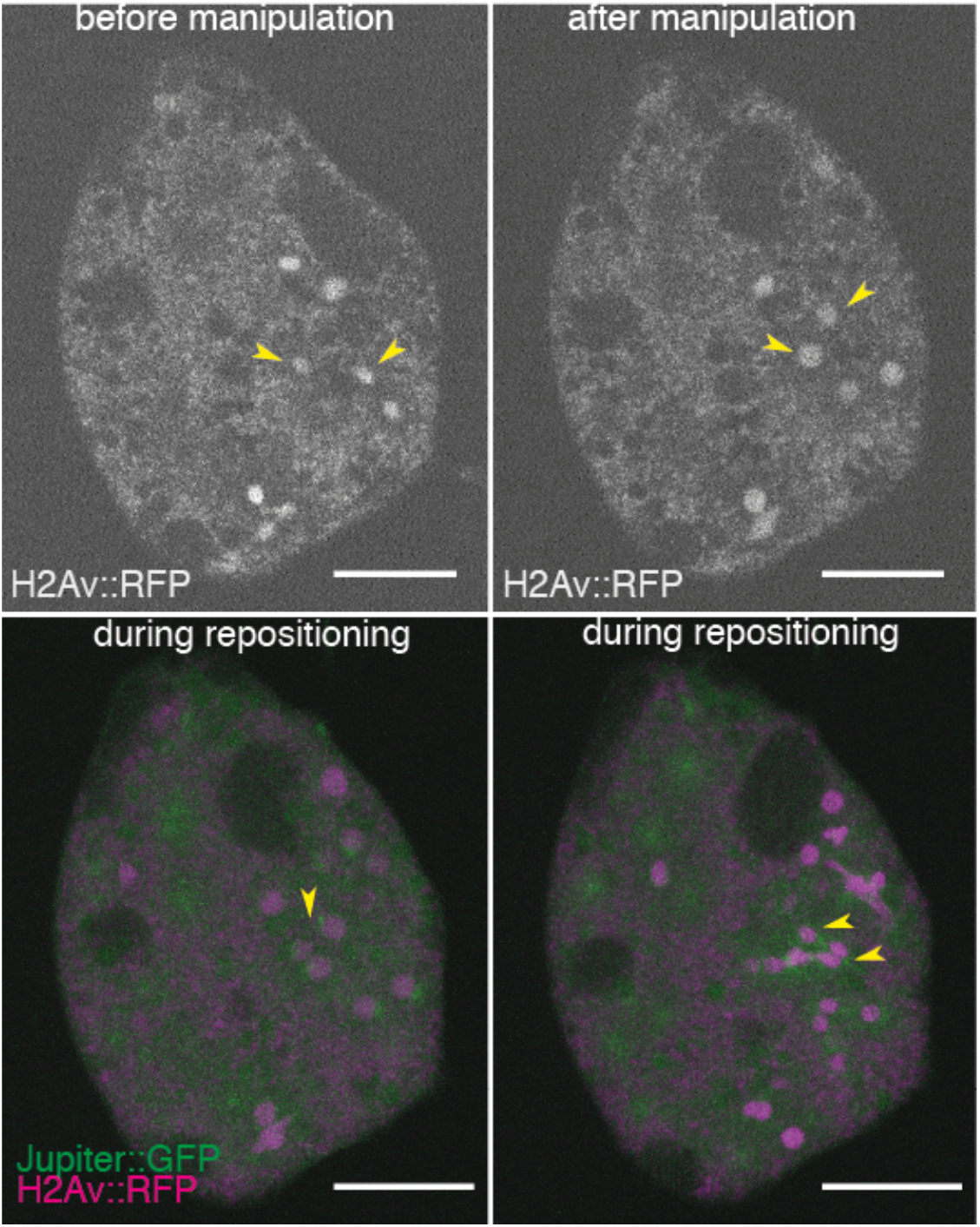
(related to Figure 4): In explants inhibited for Feo protein expression nuclear separation fails after manipulation. Still images during physical manipulation of nuclear position in an explant made from an embryo expressing RNAi against *feo* and expressing Jupiter::GFP (green) marking microtubules and H2Av::RFP (magenta) marking chromatin. After manipulation, the nuclei failed to elicit an efficient repositioning response as observed in the control (Fig. 4). Instead, sister and non-sister nuclei failed to separate sufficiently, and nuclei came into contact or formed clusters. Scale bar, 30 μm.

**Supplementary Figure 5.**
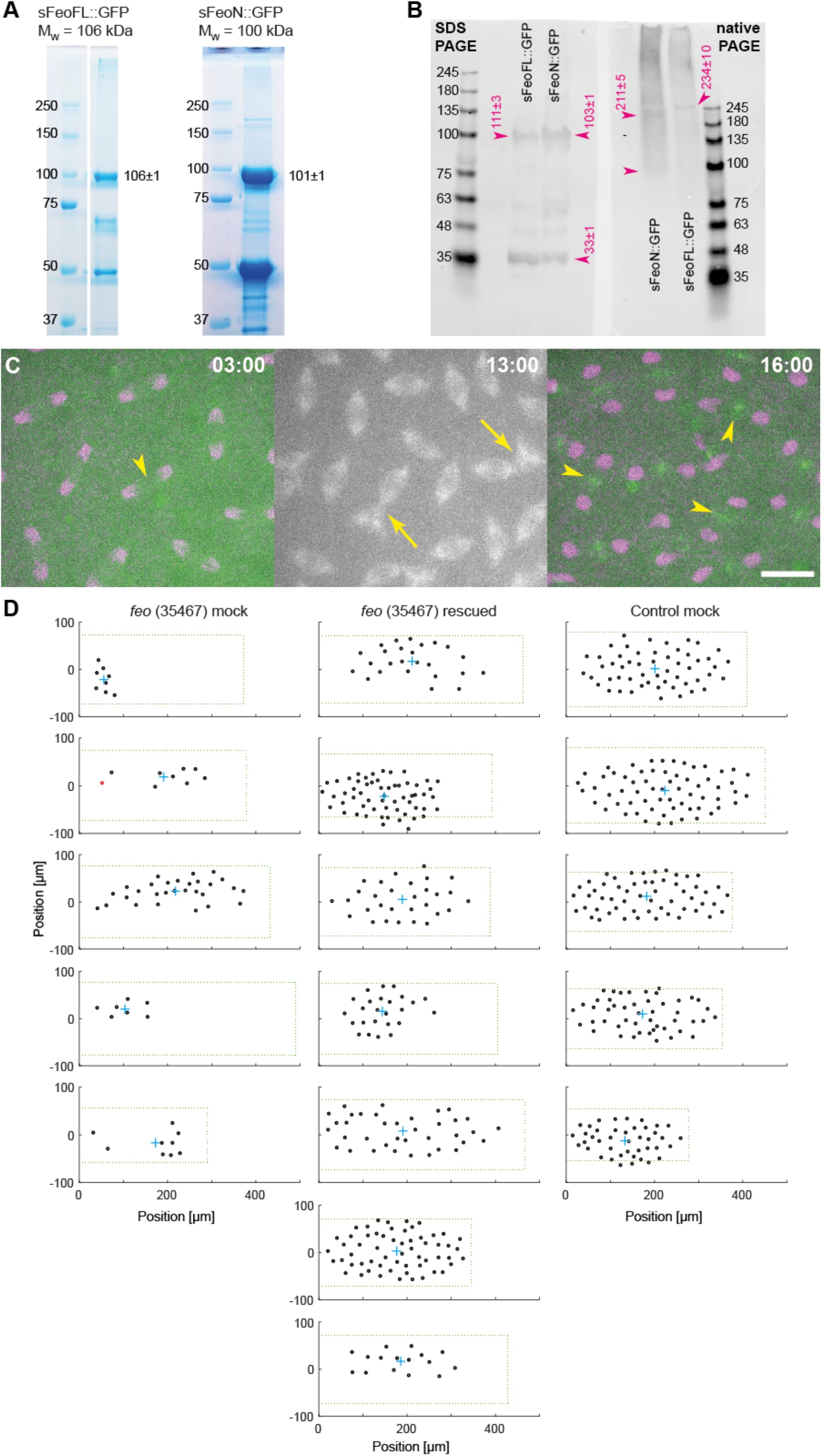
(related to Figure 5): N-terminally truncated Feo::GFP weakly dimerizes and causes nuclear separation defects, and full-length Feo::GFP protein partially rescues nuclear delivery to the cortex of *feo* RNAi embryos. **A)** Coomassie-stained 15% SDS PAGE of heterologous expressed and purified full-length Feo::GFP (left) and an N-terminally truncated Feo::GFP construct missing the putative dimerization and Klp3A-binding domain. The expected molecular mass is at the top, the measured mass is on the right of the bands (details in Methods). The lower bands are contaminants that were not separated by gel filtration and are of bacterial origin as determined by mass spectrometry. **B)** Western blot of a 4-12% gradient SDS PAGE (left) next to a Native-PAGE (right) with FeoFL:GFP and FeoN:GFP, using mouse anti-GFP antibody. Both protein constructs migrate according to expected molecular mass of a monomer in the SDS PAGE gel. In the Native-PAGE gel the full-length Feo migrates as expected from a dimer. The slight deviation from the expected mass of a dimer (212 kDa) can be explained by the rod-like structure of the protein, as was shown for the human homolog Prc1 (Subramanian et al. 2010). Conversely, the truncated construct migrates like a weak dimer, causing a wide distribution ranging between 100–200 kDa. **C)** Time-lapse fluorescence images of a (control) blastoderm embryo expressing H2Av::RFP (magenta) after injection of FeoN::GFP protein (green) and Alexa647-Tubulin (grey). During the cycle just after injection (left), the localization of FeoN::GFP at the spindle midzone was not detectable (arrowhead). However, the effect of protein addition manifests in nuclear separation defects, and in the subsequent division (middle) spindles are observed in unnatural proximity leading to spindle fusion (arrow). In telophase of the same cycle (right) FeoN::GFP is detected at the spindle midzone (arrowheads). This delayed localization after injection is not observed for FeoFL::GFP (Fig. 5B) and indicates that dimerization of Feo could be dynamic, cell-cycle dependent and, consequently, heterodimers of endogenous and truncated Feo are formed. Time is in min:sc. Scale bar, 10 μm. **D)** The position (circle) of every nucleus arriving at the embryo cortex after the last preblastoderm division, relative to the axial and lateral borders of the embryo, for each condition: *feo* (35467) mock-injected (buffer), *feo* (35467) rescued by protein injection, Control (*mcherry*) mock-injected. The green dashed rectangle represents the area of the embryo bounded by the length and width of the visible embryo in the confocal stacks, with the anterior end at the coordinate origin. The cyan cross represents the location of the 2-dimensional centroid defined from the position of all nuclei. The nuclei in interphase of the first division at the cortex are marked in black, nuclei that have progressed to metaphase / anaphase are marked in magenta.

## Supplementary Videos

**Suppl. Video 1:** Maximum intensity Z-projection from a 3D time-lapse movie of an explant generated from an embryo expressing Klp61F::GFP (cyan), Feo::mCherry (green) and injected with Alexa647-Tubulin (magenta). Time in h:min:ss, scale bar 30 μm, frame rate is 10 sec^-1^. In support of Fig. 1.

**Suppl. Video 2:** Maximum intensity Z-projection from a 3D time-lapse movie of an embryo expressing Klp61F::GFP (cyan), Feo::mCherry (green) and injected with Alexa647-Tubulin (magenta). Time in h:min:ss, scale bar 10 μm, frame rate is 10 sec^-1^. In support of Fig. 1.

**Suppl. Video 3:** Maximum intensity Z-projection from a 3D time-lapse movie of an embryo expressing Klp3A::GFP (cyan), Feo::mCherry (green) and injected with Alexa647-Tubulin (blue). Time in h:min:sec, scale bar 10 μm, frame rate is 10 sec^-1^. In the first part and 4 min^-1^ in the second part of the video. In support of Fig. 1.

**Suppl. Video 4:** Single plane time-lapse movie of an explant generated from an embryo expressing Jupiter::GFP (green) and H2Av::RFP (magenta), after addition of Nocodazole on the left side of the explant using a fine micropipette, aiming a final concentration of ∼4 μM. Time in min:sec, scale bar 50 μm, frame rate is 10 sec^-1^.

**Suppl. Video 5:** Maximum intensity Z-projection from four 3D time-lapse movies of embryos expressing Jupiter::GFP (green) and H2Av::RFP (magenta), expressing RNAi against mCherry (Control), *feo* (35467), *klp3a* (43230) and *klp61f* (30804). Time in min:sec, scale bar 50 μm, frame rate is 10 sec^-1^. In support of Fig. 2.

**Suppl. Video 6:** Maximum intensity Z-projection from four 3D time-lapse movies of explants expressing Jupiter::GFP (green) and H2Av::RFP (magenta), expressing RNAi against *mcherry* (Control), *feo* (35467) and *klp3a* (43230). Time in min:sec, scale bar 30 μm, frame rate is 10 sec^-1^. In support of Fig. 3.

**Suppl. Video 7:** Maximum intensity Z-projection from a 3D time-lapse movie of an embryo expressing Klp61F::GFP (green) and Feo::mCherry (magenta). Note that during manipulation there is additional signal from brightfield illumination, which helped visualize the glass cantilever. Time in min:sec, scale bar 30 μm, frame rate is 10 sec^-1^. In support of Fig. 4.

**Suppl. Video 8:** Maximum intensity Z-projection from a 3D time-lapse movie of an embryo expressing Spd::GFP (green, marking centrosomes), Jupiter::mCherry (magenta) and injected with sFeoFL::GFP protein (green). Time in h:min:sec, scale bar 10 μm, frame rate is 10 sec^-1^. In support of Fig. 5.

**Suppl. Video 9:** Maximum intensity Z-projection from a 3D time-lapse movie of an embryo expressing RFP::β-Tubulin (magenta) and injected with sFeoN::GFP (green) and Alexa647-Tubulin (grey). The merge shows the GFP and RFP only. Time in h:min:sec, scale bar 30 μm, Frame rate is 10 sec^-1^. In support of Fig. 5.

